# Antiviral Mechanisms and Preclinical Evaluation of Amantadine Analogs that Continue to Inhibit Influenza A Viruses with M2^S31N^-Based Drug Resistance

**DOI:** 10.1101/2024.09.09.612141

**Authors:** Ian Tietjen, Daniel C. Kwan, Annett Petrich, Roland Zell, Ivi Theodosia Antoniadou, Agni Gavriilidou, Christina Tzitzoglaki, Michail Rallis, David Fedida, Francesc X. Sureda, Cato Mestdagh, Lieve Naesens, Salvatore Chiantia, F. Brent Johnson, Antonios Kolocouris

## Abstract

To better manage seasonal and pandemic influenza infections, new drugs are needed with enhanced activity against contemporary amantadine- and rimantadine-resistant influenza A virus (IAV) strains containing the S31N variant of the viral M2 ion channel (M2^S31N^). Here we tested 36 amantadine analogs against a panel of viruses containing either M2^S31N^ or the parental, M2 S31 wild-type variant (M2^WT^). We found that several analogs, primarily those with sizeable lipophilic adducts, inhibited up to three M2^S31N^-containing viruses with activities at least 5-fold lower than rimantadine, without inhibiting M2^S31N^ proton currents or modulating endosomal pH. While M2^WT^ viruses in passaging studies rapidly gained resistance to these analogs through the established M2 mutations V27A and/or A30T, resistance development was markedly slower for M2^S31N^ viruses and did not associate with additional M2 mutations. Instead, a subset of analogs, exemplified by 2-propyl-2-adamantanamine (**38**), but not 2-(1-adamantyl)piperidine (**26**), spiro[adamantane-2,2’-pyrrolidine] (**49**), or spiro[adamantane-2,2’-piperidine] (**60**), inhibited cellular entry of infectious IAV following pre-treatment and/or H1N1 pseudovirus entry. Conversely, an overlapping subset of the most lipophilic analogs including compounds **26**, **49**, **60**, and others, disrupted viral M2-M1 protein colocalization required for intracellular viral assembly and budding. Finally, a pilot toxicity study in mice demonstrated that **38** and **49** were tolerated at doses approaching those of amantadine. Together, these results indicate that amantadine analogs act on multiple, complementary mechanisms to inhibit replication of M2^S31N^ viruses.

**Highlights:**

- Current IAVs have M2 mutations that confer resistance to amantadine and rimantadine
- Several amantadine analogs inhibit these viruses without acting on M2 proton currents
- Alternative antiviral targets include IAV entry and M2-M1 protein colocalization
- Amantadine analogs are also tolerated in mice
- Future amantadine antivirals could simultaneously act on multiple IAV mechanisms

## 1. Introduction

Despite the wide accessibility of seasonal vaccines, influenza A virus (IAV) remains a significant cause of human morbidity and mortality, contributing to approximately 0.5 million deaths annually, with the potential for many more fatalities from recurring pandemic events. As IAV has the ability to become resistant to all major classes of influenza drugs,^1,2^ additional antivirals that can target multiple drug-resistant forms of the virus are needed to support seasonal and pandemic preparedness.

The viral matrix protein 2 (M2) is required for IAV replication and is an established antiviral target. Among its multiple functions, M2 acts as a proton-gated proton channel that acidifies endosomes to promote hemagglutinin (HA)-mediated viral-host membrane fusion and release of viral RNA into the cytoplasm.^3^ During virus assembly, M2 also recruits the matrix protein 1 (M1) to the assembly site and facilitates the budding of new virions by inducing host cell membrane curvature needed for scission.^4–7^ Amantadine (*Amt*, **1**) and rimantadine (*Rmt*, **2**) (**Figure 1**) inhibit virus uncoating and assembly by acting directly on M2 proton conduction.^8–13^ Resistance to these inhibitors arises from mutations in the M2 transmembrane domain, where over 95% of adamantane-resistant IAVs bear a serine-to-asparagine substitution at position 31 (M2^S31N^), although substitutions like L26F, V27A, A30T, and G34E are also observed *in vitro* and/or in circulating viruses.^14^

**Figure 1.**
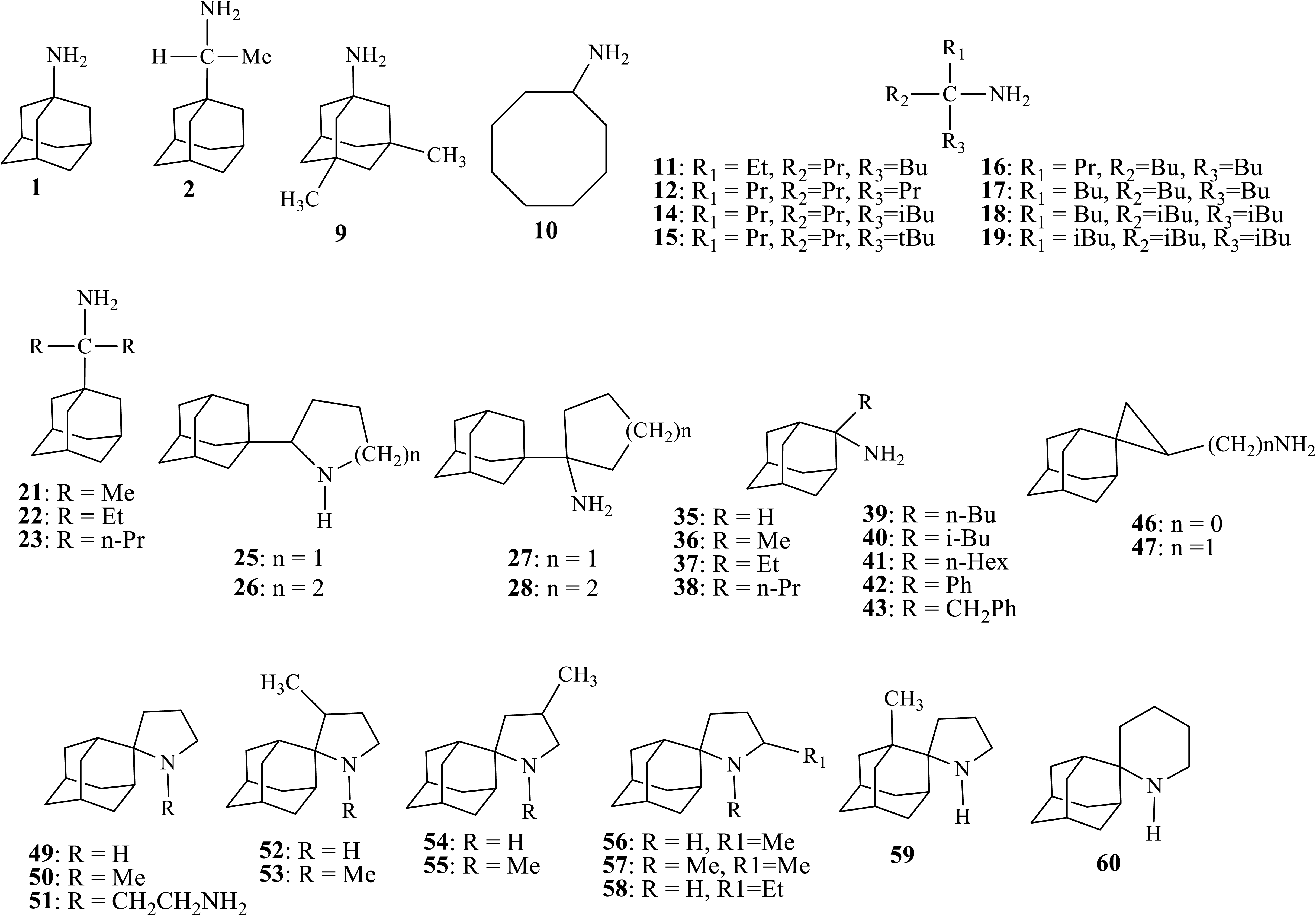
Chemical structures of *Amt*(**1**) analogs tested against M2^WT^ and M2^S31N^ IAV.

After early work identified some potent *Amt*(**1**) analogs,^15^ numerous analogs have been shown to inhibit M2^WT^ and/or *Amt*(**1**)-resistant IAVs (mostly with M2^L26F^ or M2^V27A^).^16–22^ For example, we described the antiviral activities of 57 synthetic *Amt*(**1**) analogs against (a) the drug-resistant A/H1N1/WSN/1933 virus containing M2^S31N^; (b) an A/H1N1/WSN/1933 revertant strain bearing an M2^WT^ sequence (i.e., N31S); and/or (c) A/H1N1/WSN/1933-M2^WT^ strains with additional *Amt*(**1**)-resistance markers like L26F, V27A, A30T, or G34E.^23^ Of note, several of these compounds inhibited A/H1N1/WSN/1933 viruses containing M2^WT^, M2^L26F^, and/or M2^V27A^ as well as the corresponding M2 proton currents in electrophysiology (EP) experiments. However, while none of these inhibited the unmodified A/H1N1/WSN/1933 strain bearing M2^S31N^ nor M2^S31N^-mediated proton currents,^19,22,23^ some analogs could inhibit other IAV strains containing M2^S31N^.^19,23^ For instance, we found that compounds **38** and **49** (**Figure 1**) inhibited both A/H1N1/California/07/2009, containing M2^S31N^, and A/H1N1/PR/8/1934, containing M2^A30T+S31N^, at low micromolar concentrations despite no inhibition of their M2^S31N^ proton currents.^19,23–33^

To date, the mechanisms by which *Amt*(**1**) analogs inhibit many M2^S31N^-containing viruses without directly blocking M2^S31N^ proton currents are poorly understood. For example, while some IAV inhibitors can act as lysosomotropic agents that inhibit viral-host membrane fusion by increasing endosomal pH,^24,25^ the extent to which *Amt*(**1**) analogs employ this or other antiviral mechanisms has not been investigated. To address this question, we evaluated 36 *Amt*(**1**) analogs ^23^ against a panel of IAVs containing either M2^S31N^ (A/H1N1/California/07/2009, A/H1N1/PR/8/1934, A/H1N1/WS/1933) or M2^WT^ (A/H3N2/Victoria/3/1975, A2/H2N2/Taiwan/1/1964). Selective analogs with low micromolar activity against one or more IAV with M2^S31N^ were then characterized for their additional mechanisms of action and *in vivo* tolerability.

## 2. Materials and Methods

Detailed Materials and Methods can be found in Supporting Information.

## 3. Results

### 3.1. Tested Amt(**1**) analogs

**Figure 1** shows structures of 36 assessed compounds. We tested memantine (**9**), cyclooctylamine (**10**), and **11-19** as *tert*-alkyl amine analogs of *Amt*(**1**). Compounds **21-23, 25- 28** are based on *Rmt*(**2**) with a linear alkyl, heterocyclic or carbocyclic substitution. Compounds **35-43, 46,47, 49-60** have an amino group at the C-2 position. Compounds **35-43** include an alkyl group at the C-2 adamantane carbon, **46** and **47** have a spirocyclopropyl moiety, **49-59** contain a spiropyrrolidine moiety, and **60** contains a spiropiperidine moiety.

### 3.2. Amt(**1**) analogs inhibit both M2^WT^- and M2^S31N^-containing IAVs

To elaborate on our previous observations that *Amt*(**1**) analogs such as **38** and **49** inhibit IAV without affecting the M2^S31N^ proton currents,^19,23^ *Amt*(**1**) analogs were tested against a broader range of M2^S31N^ and M2^WT^ IAVs. M2^S31N^ viruses included A/H1N1/California/07/2009, A/H1N1/PR/8/1934 and A/H1N1/WS/1933, while M2^WT^ viruses included A/H3N2/Victoria/3/1975 and A2/H2N2/Taiwan/1/1964 (**Table 1**). All compounds exhibited less than 50% cytotoxicity in MDCK cells at 50 μM or higher except for compound **27** which caused 50% cytotoxicity at 20 μM (data not shown). Using this approach, we found that *Rmt*(**2**) demonstrated strong activity against M2^WT^ viruses, with half-maximal effective concentration (EC_50_) values of 0.5 µM and 1.6 μM against A/H3N2/Victoria/3/1975 and A2/H2N2Taiwan/1/1964, respectively (**Table 1**). Consistent with previous results,^19,34^ *Rmt*(**2**) did not inhibit two M2^S31N^ viruses (EC_50_ = 106 and 314 μM against A/H1N1/California/07/2009 and A/H1N1/WS/1933, respectively), although it was active against A/H1N1/PR/8/1934 (EC_50_ = 3.3 μM) despite the presence of M2^S31N+A30T^.

**Table 1.**
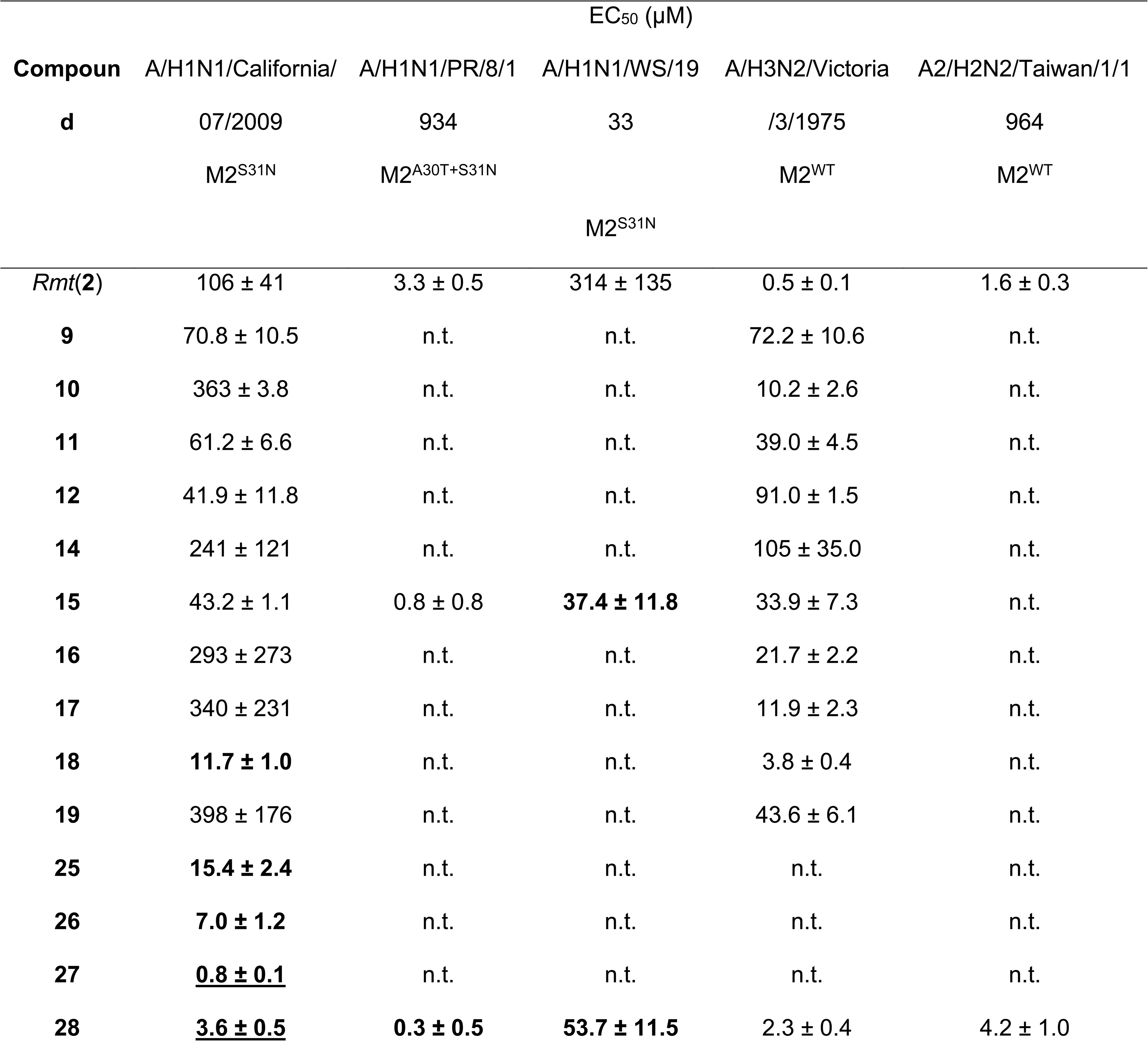
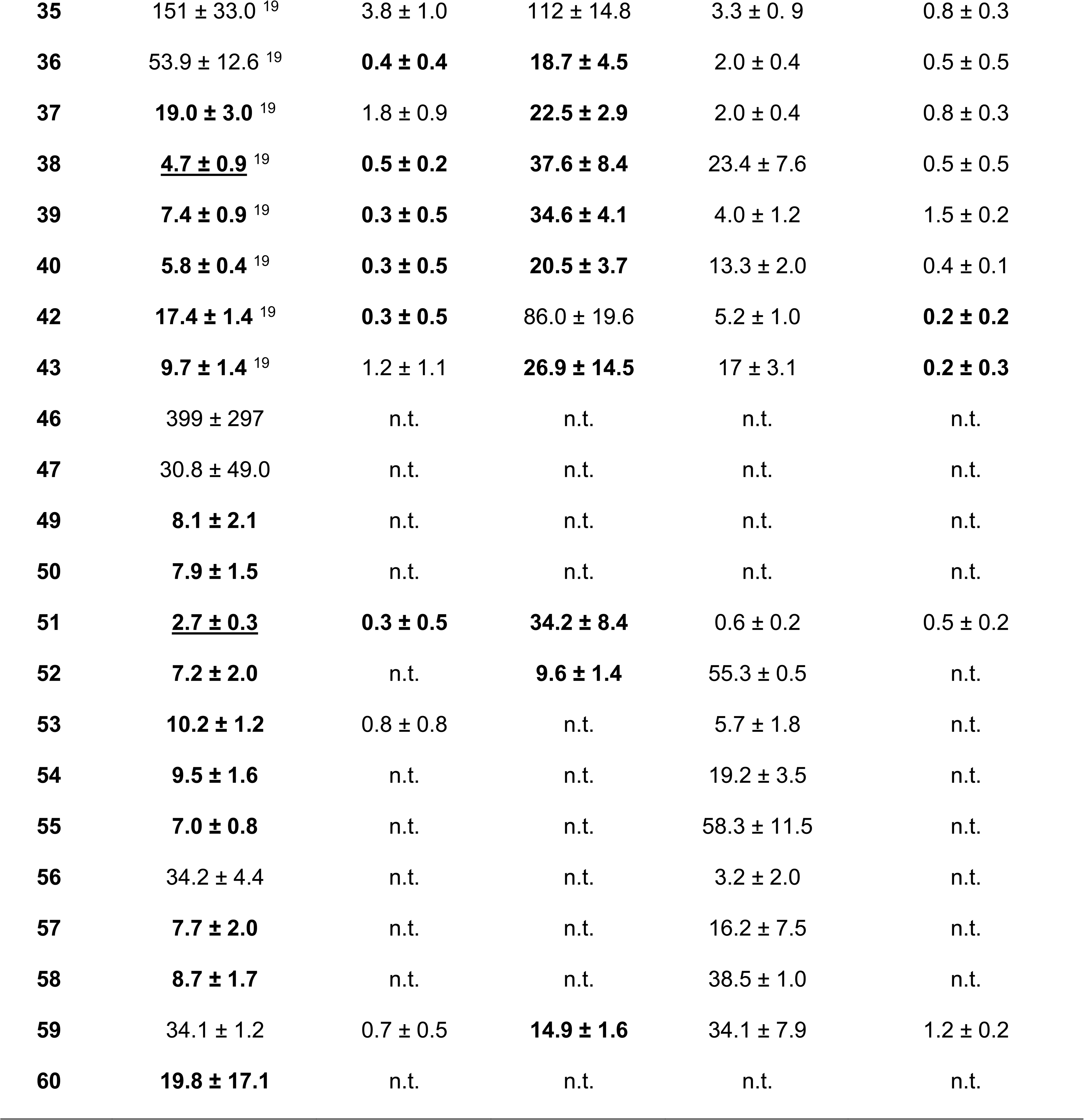
Antiviral activity and cell cytotoxicity against influenza A M2^WT^ and M2^S31N^ viruses. EC50s ± standard values were obtained from mini-plaque testing for dose-response or single-dose screens, using cultured cells, based on least-squares fitting of single-site binding curves. Values in bold reflect EC50s > 5-fold the EC50 of *Rmt*(**2**). Underlined values denote EC50s < 5 μM for M2^S31N^ viruses. n.t., not tested.

Several *Amt*(**1**) analogs were also clearly active against M2^S31N^ IAVs (**Table 1**). Against the A/H1N1/California/07/2009 virus, compounds **18**, **25-28**, **37-40, 42-43**, **49-55**, **57-58** and **60** exhibited high activity (defined here as an EC_50_ < 5 μM) or moderate activity (defined as an EC_50_ < 20 μM or ∼5-fold lower than *Rmt*(**2**)), with **27** being the most active (EC_50_ = 0.8 μM, or 132-fold lower than *Rmt*(**2**)). Moderate activity against A/H1N1/PR/8/1934 (*i.e.,* EC_50_ < 0.65 μM or 5-fold lower than *Rmt*(**2**)) was observed for compounds **28, 36, 38-40, 42** and **51**. When assessed against A/H1N1/WS/1933, compounds with EC_50_ < 65 μM (*i.e.,* 5-fold lower than *Rmt*(**2**)) included **15, 28, 36-40, 43, 51-52** and **59**. Overall, five compounds (**28, 38-40** and **51**) exhibited EC_50_ values at least 5-fold lower than *Rmt*(**2**) across all three M2^S31N^ viruses (**Table 1**). Previous studies using EP showed that compounds **28** and **38** (as well as **35** and **49**) do not block proton currents of M2^S31N^ from A/H1N1/California/07/2009.^35^

In contrast, several *Amt*(**1**) analogs exhibited detectable activity (defined here as EC_50_ < 12.5 μM or within 25-fold of *Rmt*(**2**)) against the A/H3N2/Victoria/3/1975 IAV strain with M2^WT^; these included **10**, **17-18**, **28**, **35-37**, **39**, **42**, **51**, **53**, and **56** (**Table 1**). Compounds **28**, **35-40**, **42-43**, **51**, and **59** also showed detectable activity against the A2/H2N2/Taiwan/1/1964 virus (*i.e.*, EC_50_ < 12.5 μM or within 25-fold of *Rmt*(**2**)) (**Table 1**). Consistent with these observations, compounds **28, 38,** and **51** inhibit proton currents from the M2^N31S^-mutated proton channel of A/H1N1/California/07/2009 (*i.e.,* reverted to WT).^35^ However, contrary to observations with M2^S31N^ viruses, where multiple analogs had at least 5-fold improvement over *Rmt*(**2**), no analog had this level of improvement over *Rmt*(**2**) except for compounds **42** and **43** against A2/H2N2/Taiwan/1964 (**Table 1**).

To determine whether *Amt*(**1**) analog activities were consistent across M2^S31N^ and M2^WT^ viruses, we performed linear regression analysis of the average EC_50_ for each compound across viruses (**Figure 2**). Notably, the EC_50_ values of all compounds tested against A/H1N1/California/07/2009 and A/H1N1/PR/8/1934 which both contain M2^S31N^, were well correlated (r^2^ = 0.77; p < 0.0001; **Figure 2A**). Significant correlations were also maintained between the EC_50_s of A/H1N1/California/07/2009 and A/H1N1/PR/8/1934 versus A/H1N1/WS/1933, which also has M2^S31N^ (r^2^ = 0.41; p = 0.013 and 0.47; p = 0.009, respectively; **Figure 2B-C**). These results suggest that *Amt*(**1**) analogs may act on the same viral target(s) across these M2^S31N^ viruses. In contrast, no correlations were observed between M2^S31N^ and M2^WT^ viruses (all r^2^ < 0.13; p > 0.05; **Figure 2D**), suggesting that the primary viral target(s) differ between these two groups. Similarly, no correlation was found between A/H3N2/Victoria/3/1975 and A2/H2N2/Taiwan/1/1964 (both M2^WT^) (**Figure 2E**).

**Figure 2.**
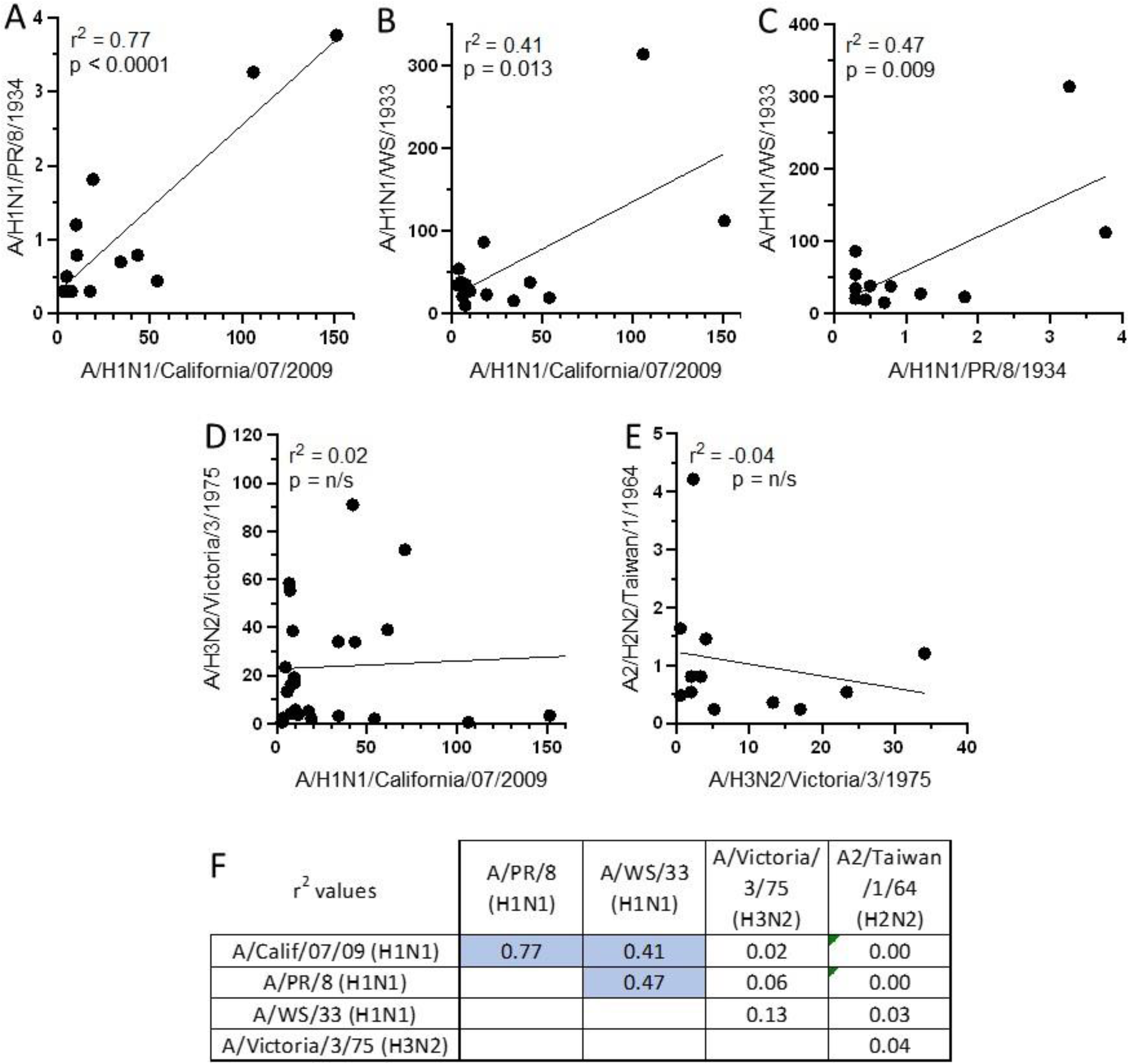
Comparisons of compound EC_50_s (in µM) across viruses in **Table 1**. **A-E,** Representative comparisons of EC_50_s of A/H1N1/PR/8/1934 vs. A/H1N1/California/07/2009 (**A**), A/H1N1/WS/1933 vs. A/H1N1/California/07/2009 (**B**), A/H1N1/WS/1933 vs. A/H1N1/PR/8/1934 (**C**), A/H3N2/Victoria/3/1975 vs. A/H1N1/California/07/2009 (**D**), and A2/H2N2/Taiwan/1/1964 vs. A/H3N2/Victoria/3/1975 (**E**). **F,** Summary of all correlations. For each comparison, at least 12 compounds were assessed including *Rmt*(**2**). Blue shading denotes r^2^ values with a linear regression p-value < 0.05. n/s, non-significant.

### 3.3. Antiviral activity of Amt(**1**) analogs is not due to inhibition of M2^S31N^-dependent proton currents

To rule out any direct effect on M2^S31N^ proton conduction, we used a previously- described whole-cell patch-clamp EP approach ^27^ with compound **38** representing analogs that inhibited all three M2^S31N^ viruses. Briefly, tsA-201 cells co-transfected with plasmids encoding GFP and M2^S31N^ (from A/H1N1/California/07/2009) were held at a constant membrane potential of -40 mV, and currents were recorded every 4 s by applying 100-ms pulses to -80 mV. Using this approach, we previously showed that robust, pH-dependent inward currents are readily inhibited with M2^WT^ and M2^S31N^-dependent inhibitors.^27^ We then assessed the activity of **38** over a 30-minute exposure period. As shown in **Figure 3A**, single tsA-201 cells expressing M2^S31N^ exhibited robust, low pH-dependent negative or inward current that decayed ∼50% during continued exposure to low pH. This current could be maintained for more than 2400 seconds (40 minutes) and was inhibited approximately 25% with 100 μM *Amt*(**1**), consistent with previous results of its fast but limited inhibition.^27^ However, when the same long-term procedure was done with 100 μM of **38**, no inhibition of M2^S31N^ current was observed even after 2400 seconds (**Figure 3B**). The finding that compound **38** does not inhibit M2^S31N^ currents even after prolonged exposure further argues against M2^S31N^ channel inhibition as the mechanism underlying the activity of compound **38** and related compounds towards M2^S31N^ viruses.

**Figure 3.**
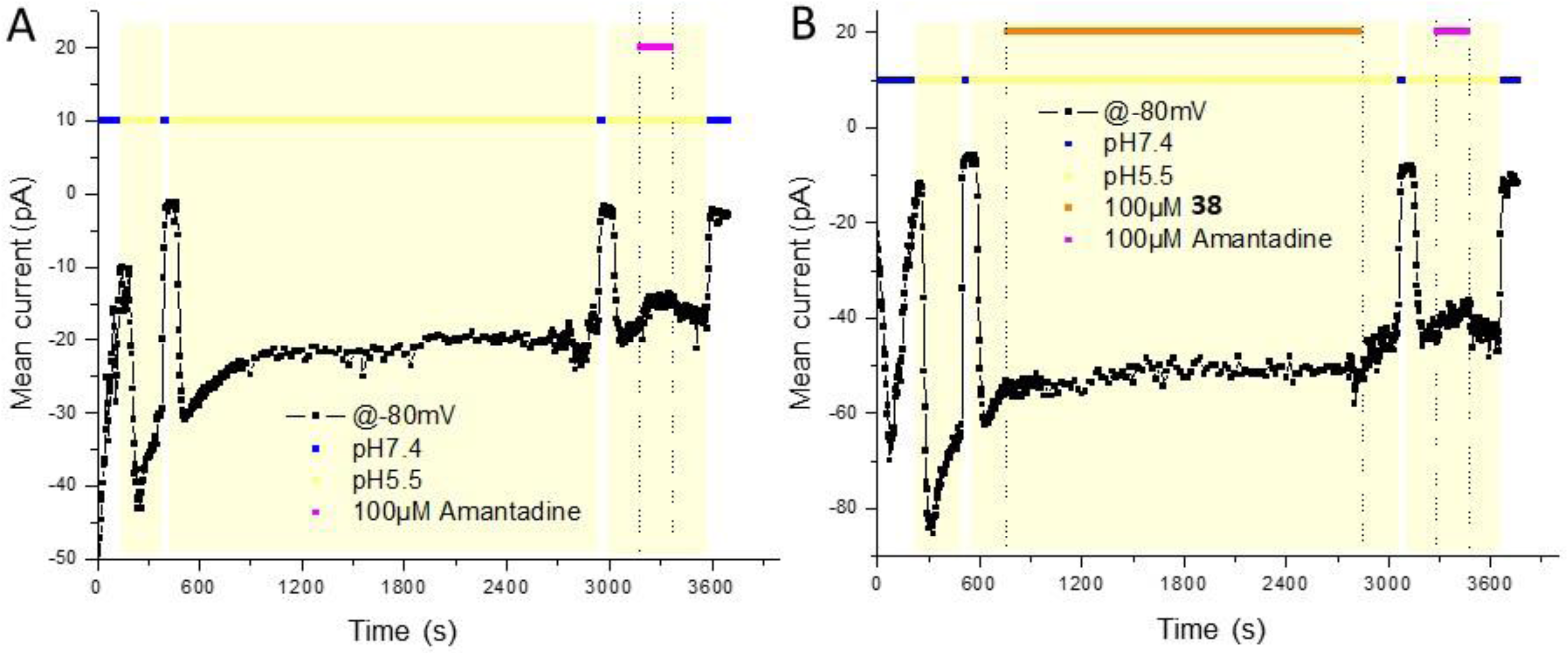
Representative diary plots of pH-dependent currents detected at -80 mV from single tsA-201 cells transiently expressing GFP and M2^S31N^ (A/H1N1/California/07/2009). Dots denote currents recorded at -80 mV every 4 seconds. Colors denote extracellular pH and addition of compounds. Effects of *Amt*(**1**) (**A**) and compound **38** (**B**) are shown.

### 3.4. Antiviral activity of Amt(**1**) analogs is not due to modulating endosomal pH

In addition to inhibiting M2, some *Amt*(**1**) analogs can function as lysosomotropic agents, preventing viral-host membrane fusion by increasing endosomal pH.^24,25^ Besides IAV, viruses like bovine papillomavirus (BPV) also rely on acidic endosomes for entry or uncoating within host cells. Compounds that elevate endosomal pH or disrupt the endocytosis process can inhibit BPV replication.^36^ To explore whether *Amt*(**1**) analogs might modulate endosomal pH, we assessed compounds **27** and **38** in a miniplaque assay using BPV (P6 variant)-infected bovine embryonic kidney cells. While control agents chlorpromazine, ammonium chloride, chloroquine and bafilomycin A1 all inhibited BPV replication at concentrations consistent with published results (EC_50_: 2.5, 0.4, 2.2 and 0.06 μM, respectively),^36^ no inhibition was observed by up to 20 μM of either *Amt*(**1**) analog (**Figure 4**), suggesting that **27** and **38**, and presumably other *Amt*(**1**) analogs, do not act as endosome neutralizers to inhibit M2^S31N^ viruses.

**Figure 4.**
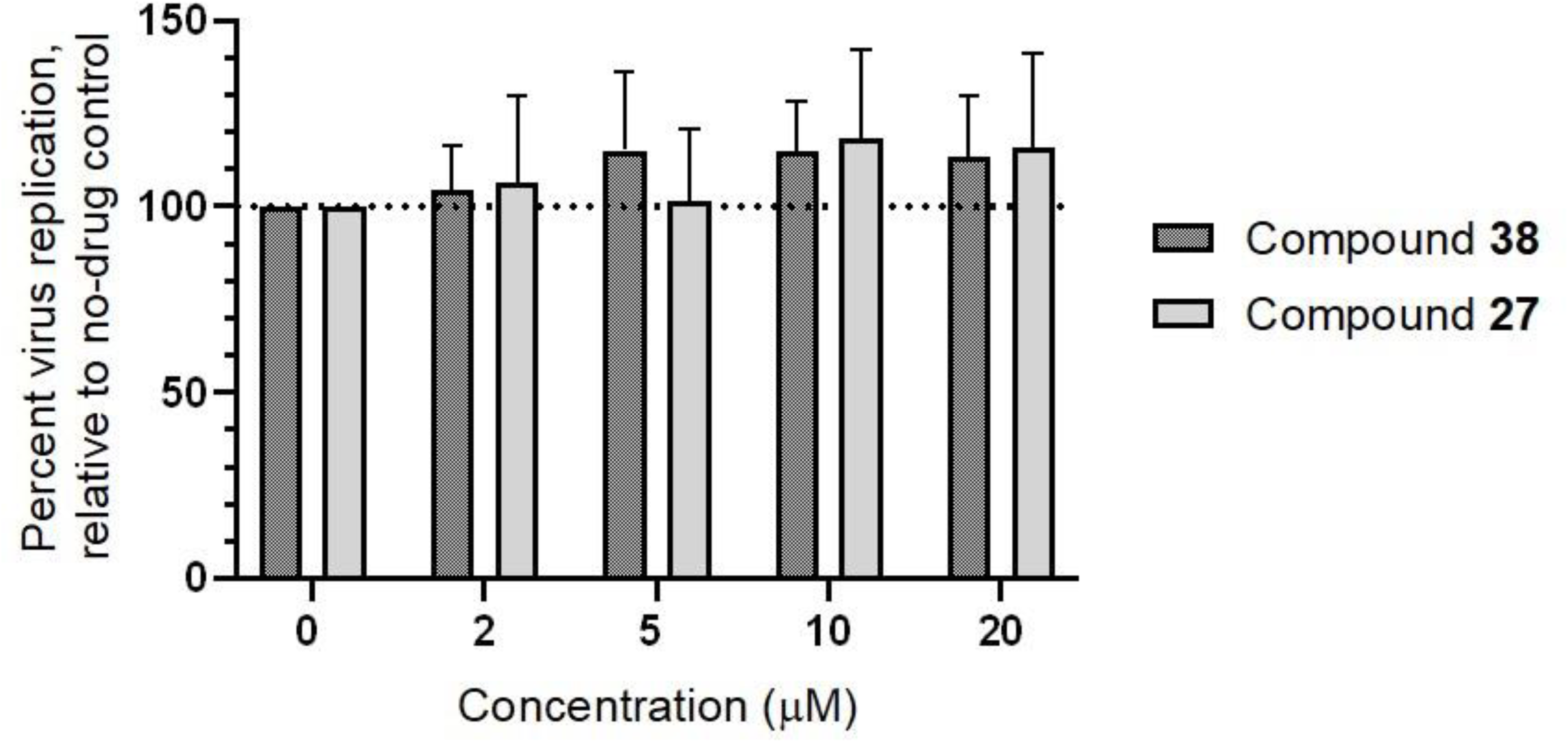
*Amt*(**1**) analogs do not inhibit BPV replication, in contrast to control antivirals that selectively act through increasing endosomal pH. Results denote mean ± SD of four independent experiments.

### 3.5. In vitro selection of Amt(**1**) analog-resistant viruses occurs through M2 mutations in M2^WT^ but not M2^S31N^ viruses

To identify potential viral targets of these *Amt*(**1**) analogs, we conducted virus passaging to select for compound-resistant viruses. MDCK cells were initially infected with A/H3N2/Hong Kong/1/1968, which contains M2^WT^, and passaged semi-weekly in the presence of 3.9, 5.0, or 4.4 μM of compounds **27**, **36**, or **38**, respectively (corresponding to 1, 1, and 5 µg/mL). As anticipated, all compounds induced rapid drug resistance. Following plaque purification and M- sequencing, resistance was linked to known *Amt*(**1**)-resistance mutations in M2 including V27A for compound **27** and A30T for **36** and **38** (**Table 2**). These results support that *Amt*(**1**) analogs readily select for escape mutants bearing substitutions in M2, suggesting that, similar to *Amt*(**1**), these analogs inhibit M2^WT^ viruses by targeting M2 proton conduction.

**Table 2.** Mutations developed influenza M2^WT^ after passaging experiments with compounds.

| A/H3N2/Hong |  |  |  |  |
| --- | --- | --- | --- | --- |
| Kong/1/1968 | Compound | 27 | 36 | 38 |
| (M2<sup>WT</sup>) |  |  |  |  |
| Passage | Plaque |  |  |  |
| number | number <sup>a</sup> | 3.9 μM | 5.0 μM | 4.4 μM |
| 2 | 1 | V27A | WT | A30T |
|  | 2 | V27A | WT | A30T |
|  | 3 | V27A | WT | A30T |
| 5 | 1 | V27A | A30T | A30T |
|  | 2 | V27A | A30V | A30T |
|  | 3 | V27A | A30T | A30T |
| 10 |  |  | 2 μg/ml | 5 μg/ml |
|  | 1 |  | A30T | A30T |
|  | 2 |  | A30T | A30T |
|  | 3 |  | A30T | A30T |

<sup>a</sup> Three separate plaques were sampled and sequenced at passages 2, 5, and/or 10.

We next used a subset of *Amt*(**1**) analogs for passaging experiments with an M2^S31N^ virus. MDCK cells infected with A/H1N1/California/07/2009 were passaged semi-weekly in the presence of 5 μM of compound **38** or a triple combination of compounds **26**, **27** and **60**, each at 5 μM, representing a diverse set of ring adducts at both C1 and C2 (**Table 3**). As an experimental control, we performed parallel passaging of A/H3N2/Victoria/3/1975 virus with M2^WT^ in the presence of 50 µM *Amt*(**1**). In this control, drug resistance emerged after one passage: while the parent virus had an EC_50_ value of 3.0 µM (n = 9) for *Amt*(**1**), passage #1 and #2 viruses were not inhibited by *Amt*(**1**). In contrast, for A/H1N1/California/07/2009 grown under compound **38**, passages #1 to #5 remained fully sensitive (EC_50_ = 2.1 – 5.4 µM), but between passages #6 and #12, the virus became progressively resistant, with > 20-fold resistance by passage #10 onward (EC_50_ = 76 – 149 µM; **Table 3**). Similarly, the combination of **26**, **27** and **60** remained effective through 6 passages (EC_50_ = 1.0 – 1.2 µM) but developed ∼6.6-fold resistance by passage #10 (EC_50_ = 7.9 µM; **Table 3**). Notably, the passage #12 virus resistant to compound **38** remained sensitive to compound **28,** with an EC_50_ of 10.2 µM only modestly increased compared to the parental virus (EC_50_ = 3.6 µM). Sequence analysis on the parental A/H1N1/California/07/2009 virus and passage #12 virus resistant to compound **38** indicated no differences in the region encoding M2 amino acids 10-73. Hence, resistance to **38** was not caused by additional mutations in M2 that underlie proton conduction.

**Table 3.** Resistance testing of amantadine (1) variants against an amantadine-sensitive H3N2 virus and potent amantadine variants against amantadine-resistant H1N1 (2009).

|  | **1** (5 µM) | **38** (5 µM) | **26, 27, 60** (5 µM) |
| --- | --- | --- | --- |
|  | A/H3N2/Victoria/3/1975 | A/H1N1/California/07/2009 | A/H1N1/California/07/2009 |
| Passage # | (M2<sup>WT</sup>) | (M2<sup>S31N</sup>) | (M2<sup>S31N</sup>) |
|  | EC<sub>50</sub> ± S.E. (µM) | EC<sub>50</sub> ± S.E. (µM) | EC<sub>50</sub> ± S.E. (µM) |
| 0 | 2.8 ± 0.3 | 4.7 ± 0.9 | 1.0 ± 0.2 |
| 1 | > 50 | 5.4 ± 1.4 |  |
| 2 | > 50 | 3.7 ± 0.5 |  |
| 5 |  | 2.1 ± 1.6 |  |
| 6 |  |  | 1.2 ± 0.1 |
| 8 |  | 18.5 ± 1.0 |  |
| 10 |  | 76 ± 9 | 7.9 ± 0.8 |
| 12 |  | 149 ± 115 |  |
| Compound |  | 10.2 ±1.7 |  |
| **28** |  |  |  |

Taken together, these results indicate *Amt*(**1**) analogs primarily act on M2 in viruses containing M2^WT^ but not M2^S31N^. Additionally, since mutations selected during passaging viruses containing M2^S31N^ did not confer significant resistance across all *Amt*(**1**) analogs, it is likely that these analogs act on multiple and potentially overlapping viral targets.

### 3.6. Amt(**1**) analogs inhibit the entry of M2^S31N^ virus in an HA-dependent manner

To determine whether *Amt*(**1**) analogs interfere with IAV’s ability to infect MDCK cells, we treated A/H1N1/California/07/2009 virus with or without 50 μM compound **38**. Virus-drug aliquots were collected at 0, 1, 5-, 10-, 20- or 30-minute post-incubation and assessed for infectivity by miniplaque assay. At this stage, the final concentration of **38** was 0.5 μM, a concentration unlikely to inhibit virus growth (**Table 1**). After 5 minutes of incubation, IAV infectivity was reduced by 46.7%; by 30 minutes, reduction reached 62.9% (**Figure 5**). These results suggest that **38** reduces virus infectivity before exposure to target cells.

**Figure 5.**
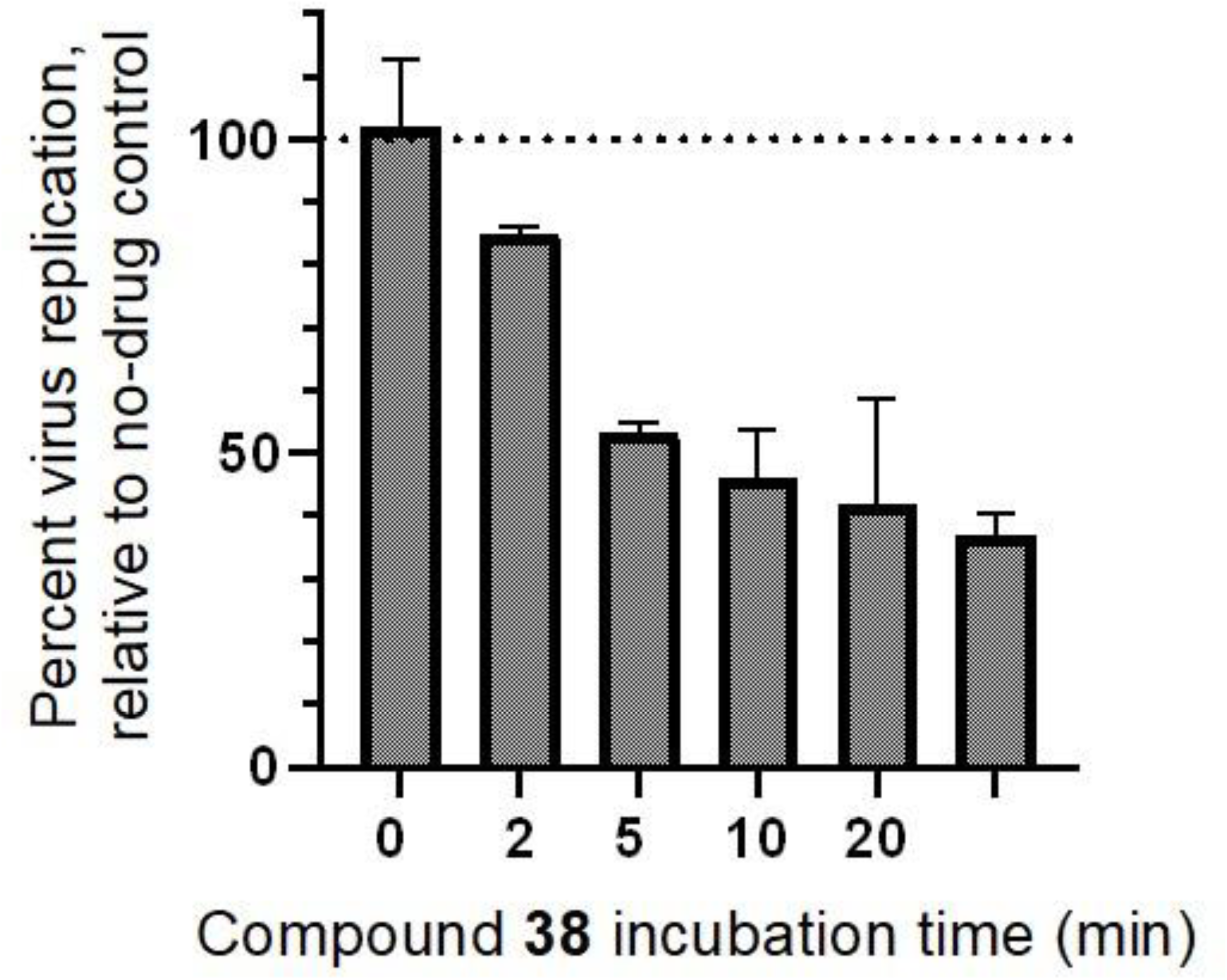
Compound **38** inhibits cellular entry of influenza virus with M2^S31N^. A/H1N1/California/07/2009 virus was incubated with 50 μM of compound **38** at defined time intervals before infection of MDCK cells. Results denote mean ± SD from two independent experiments.

To test whether **38** might act on HA-mediated viral entry,^37^ we next conducted an MDCK- based entry assay using murine leukemia virus-based pseudovirus bearing the HA and neuraminidase proteins of influenza A/Virginia/ATCC3/2009. As shown in **Figure 6**, *Amt*(**1**), *Rmt*(**2**), and compounds **26**, **49**, and **60** did not inhibit pseudovirus entry at 400 µM. In contrast, compound **38** exhibited clear activity, with an EC_50_ of 16 µM, followed by **39** and **28** with EC_50_s of 29 and 64 µM, respectively. The EC_50_ of compound **38** was comparable to those of control viral entry inhibitors arbidol and hydroxychloroquine (**Figure 6**).

**Figure 6.**
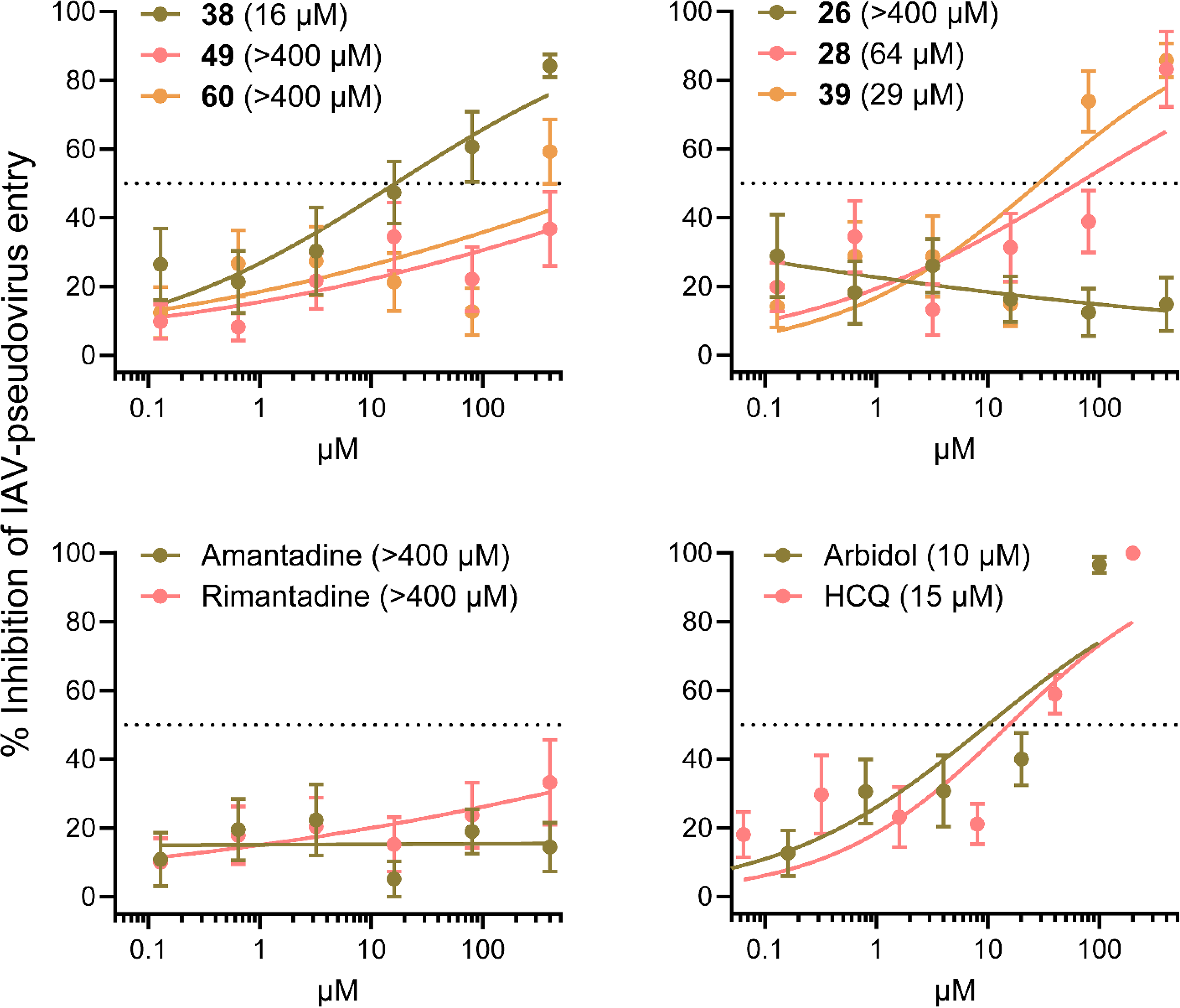
Inhibition of H1N1 pseudovirus entry by a selection of *Amt*(**1**) analogs. Pseudovirus bearing the HA and neuraminidase proteins from A/H1N1/Virginia/ATCC3/2009 virus was transduced into MDCK cells in the presence of the compounds. After three days incubation, the luciferase readout was conducted. In each panel, the EC_50_ values are shown in brackets. Each data point shows the mean ± SEM of nine values (three independent experiments, each performed in triplicate). HCQ: hydroxychloroquine.

### 3.7. Amt(**1**) analogs impair M2-mediated M1 recruitment to the plasma membrane

The compounds evaluated above (**26, 28, 38, 39, 49, 60**) were subsequently investigated for their ability to disrupt the intracellular distribution of the viral matrix proteins M1 and M2. HEK-293T cells were transfected with fluorescent constructs labelled with mEGFP and mCherry2, respectively, and their subcellular localization was monitored by fluorescence microscopy. We previously showed that M2 is essential for the proper recruitment of M1 to the plasma membrane (PM).^31^ In cells lacking M2 expression (i.e., negative control, NC in **Figure 7**), M1 was uniformly distributed throughout the cytoplasm. Conversely, in the presence of M2 (positive control, PC in **Figure 7**), M1 was enriched at the PM, as expected.^54^ Of note, treatment with 30 µM of *Amt*(**1**) analogs like compound **39** significantly decreased the number of cells with M1-PM association (**Figure 7; Figure S1**). Quantitative analysis indicated that all tested compounds significantly impaired M2-M1 interactions except for **38**, which only showed an approximately 10% reduction (**Figure 8**, **Table S1 and Figures S2-S3**). For instance, compound **39** reduced M2-M1 colocalization by approximately 63% (**Figures 7-8**).

**Figure 7.**
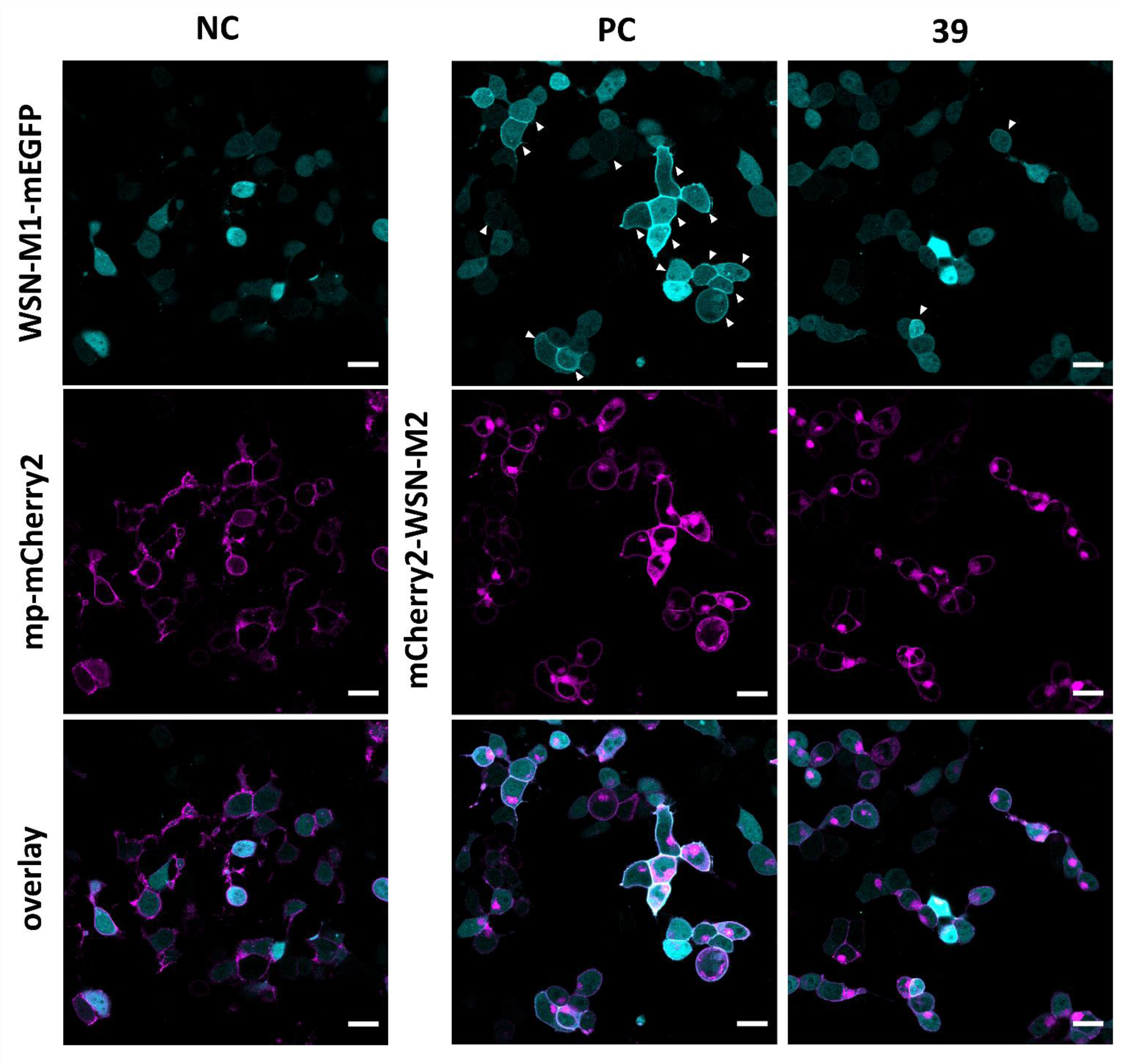
Membrane recruitment of influenza A virus matrix protein 1 (M1) in matrix protein 2 (M2) co-transfected cells is altered by *Amt*(**1**) analogs. Representative confocal fluorescence images of HEK-293T cells co-expressing WSN-M1-mEGFP (cyan) (from A/H1N1/WSN/1933) and either plasma membrane-associated mCherry2 (mp-mCherry2, as negative control (NC)), or HA_sp_-mCherry2-WSN-M2 (magenta). The latter samples were treated either with H_2_O (as positive control (PC)), or 30 µM of compound **39**. Imaging acquisition took place after 24 hours. The lower panels show the two channels merged into a single image. Images for compounds **38**, **49**, **60**, **26** and **28** at 30 µM are shown in the supporting material (**Figure S1**). White arrows indicate cells with M1 plasma membrane (PM) localization. Intensity ranges were calibrated 2 to 100 for WSN-M1-mEGFP and HA_sp_-mCherry2-WSN-M2. Scale bars represent 20 µm.

**Figure 8.**
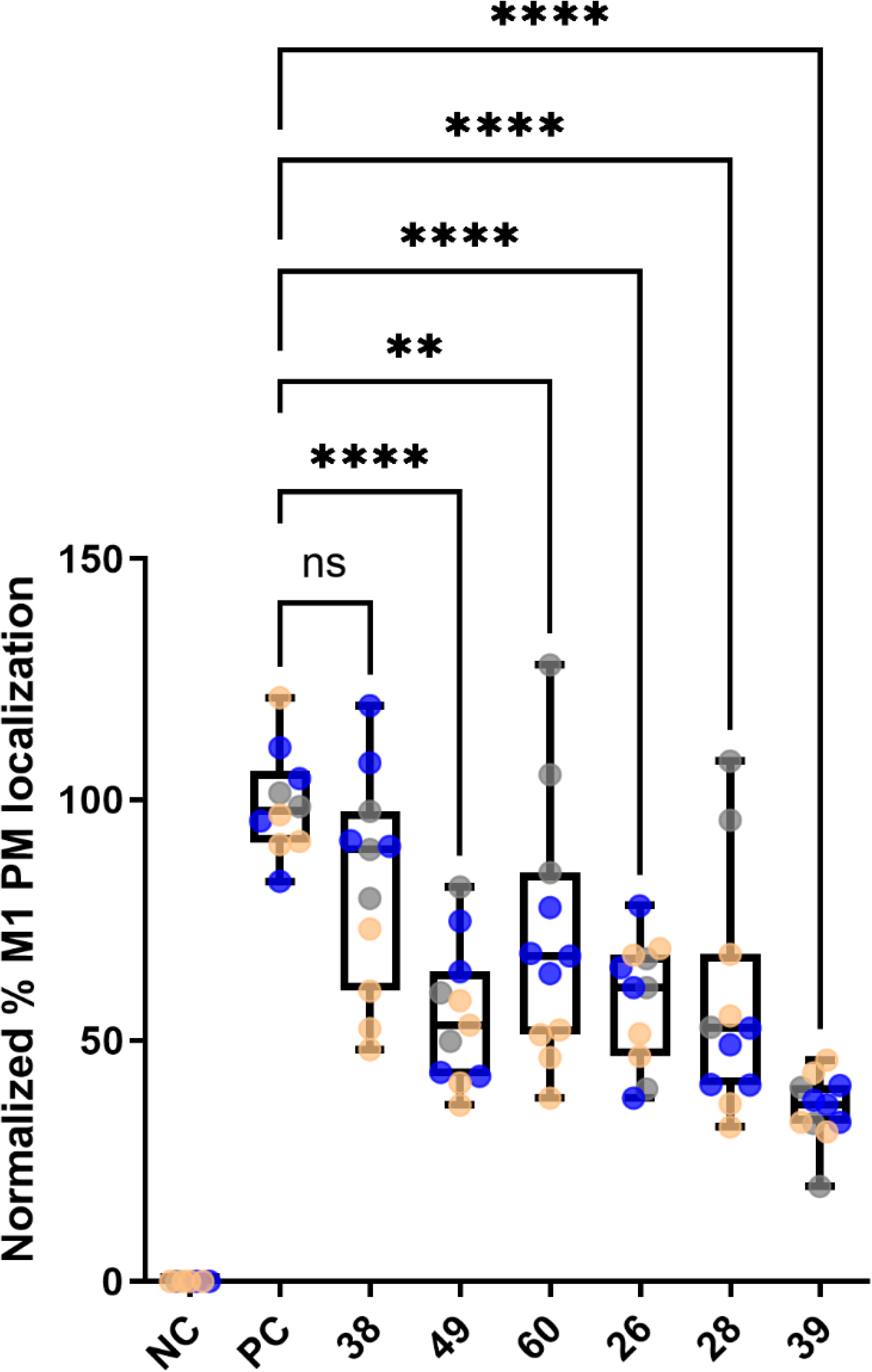
Quantification of IAV M1 recruitment to the PM by M2 in the presence of *Amt*(1) analogs. The plot shows the normalized percentage of cells displaying PM localization of M1 for the indicated treatment. A concentration of 30 µM was used for all compounds and imaged after 24 hours. The percentage of cells displaying M1 signal at the PM was manually identified using Fiji ImageJ software and the ratio of these cells to the total number of cells was calculated. This value was then normalized to the median value of the positive control (PC). Data from three separate experiments were pooled, plotted, and analyzed using a one-way ANOVA Dunnett’s multiple comparison test (** p < 0.01, **** p < 0.0001). Each data point represents the percentage value measured for one image and the colors of each point represent an individual experiment (experiment 1 (grey), 2 (blue) and 3 (orange)). Descriptive statistics are summarized in the supporting material (**Table S1**).

Based on these mechanistic insights, the overall antiviral activity of *Amt*(**1**) analogs against M2^S31N^ viruses arises from a combination of inhibitory effects on viral entry and disruption of M2-M1 colocalization.

### 3.8. Amt(**1**) analogs and combinations are tolerated in vivo

The ability of *Amt*(**1**) analogs to act on multiple aspects of IAV replication positions them as promising lead compounds for developing antivirals. These antivirals, in principle, could combine a direct effect on the M2^WT^ channel with additional effects on viral entry and/or M2-M1 colocalization in M2^S31N^-containing viruses. To evaluate *in vivo* safety, we conducted a pilot study comparing compounds **38** and **49** to *Amt*(**1**) for *in vivo* safety upon *i.p.* injection in CD-1 mice. All three compounds were tolerated at 30 mg/kg. However, compound **38** was uniformly lethal at 100 mg/kg, whereas **49** and *Amt*(**1**) were lethal at 300 mg/kg (**Table 4**). Furthermore, while **49** exhibited some signs of neurotoxicity at 30 mg/kg, no abnormalities were observed with either *Amt*(**1**) or **38** (**Table 4**). Hence, compound **38** is tolerated *in vivo* at least at 30 mg/kg and warrants further investigation for *in vivo* efficacy.

**Table 4.**
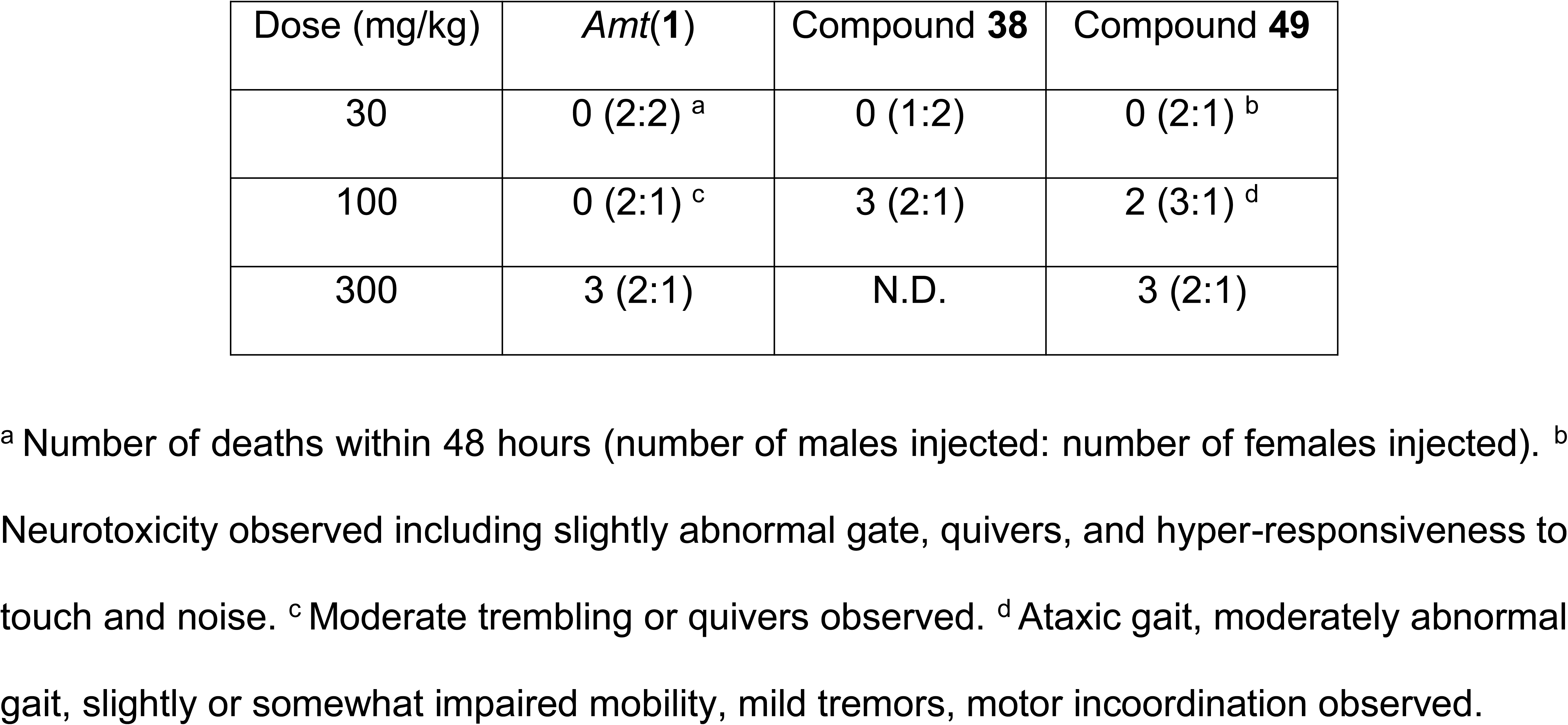
*In vivo* toxicity from intraperitoneal infections of *Amt*(1) and compounds 38 and 49.

| Dose (mg/kg) | *Amt(1)* | Compound **38** | Compound **49** |
| --- | --- | --- | --- |
| 30 | 0 (2:2) <sup>a</sup> | 0 (1:2) | 0 (2:1) <sup>b</sup> |
| 100 | 0 (2:1) <sup>c</sup> | 3 (2:1) | 2 (3:1) <sup>d</sup> |
| 300 | 3 (2:1) | N.D. | 3 (2:1) |

<sup>a</sup> Number of deaths within 48 hours (number of males injected: number of females injected). <sup>b</sup> Neurotoxicity observed including slightly abnormal gate, quivers, and hyper-responsiveness to touch and noise. <sup>c</sup> Moderate trembling or quivers observed. <sup>d</sup> Ataxic gait, moderately abnormal gait, slightly or somewhat impaired mobility, mild tremors, motor incoordination observed.

## 4. Discussion

To effectively manage seasonal and pandemic IAV outbreaks and support vaccination efforts,^1,2,3^ new antivirals are essential, particularly those capable of inhibiting drug-resistant IAV strains. Drugs that target IAV through multiple mechanisms are preferable, as they are likely to provide a higher genetic barrier to resistance. Toward this goal, we evaluated the activities of 36 previously synthesized *Amt*(**1**) analogs against a panel of IAV strains, including both M2^WT^ or *Amt*(**1**)-resistant M2^S31N^. Notably, several compounds, including **15**, **18**, **25-28**, **36-43**, **49-55**, and **57-60**, inhibited up to three M2^S31N^-containing viruses.

The activities of analogs across M2^S31N^-containing viruses were well correlated (**Figure 2**), suggesting that they share at least some antiviral mechanisms. Using various molecular techniques, we ruled out and identified antiviral targets for a subset of analogs. As shown in **Table 5**, analogs had additional properties including ability to inhibit virus entry, achieved through inhibiting virus infectivity during pre-incubation before exposure to host cells and/or through blocking H1N1 pseudovirus entry, and/or disrupting intracellular colocalization of M2 and M1, which is required for viral assembly and budding.

**Table 5.** Summary of bioactivities of select *Amt*(1) analogs. *, compounds passaged in combination.

| Compound | Inhibits M2<sup>S31N</sup> by EP at 100 μM | Inhibits BVP replication (lysosomotropic effect) at 20 μM | Mutations in M2<sup>WT</sup> develop at 1-5 μg/mL | Mutations in M2<sup>S31N</sup> develop at 5 μM | Inhibits cellular entry of IAV with M2<sup>S31N</sup> at 5 μM | H1N1 pseudo virus entry (EC<sub>50</sub>, μM) | Inhibits M1-M2<sup>S31N</sup> colocalization at 30 μM | Tolerated *in vivo* at 30 mg/kg |
| --- | --- | --- | --- | --- | --- | --- | --- | --- |
| **26** |  |  |  | No* |  | > 400 | Yes |  |
| **27** |  | No | V27A | No* |  |  | Yes |  |
| **28** | No<sup>31</sup> |  |  |  |  | 64 | Yes |  |
| **38** | No | No | A30T | No | Yes, at 50 μM | 16 | No | Yes |
| **39** |  |  |  |  |  | 29 | Yes |  |
| **49** | No<sup>31</sup> |  |  |  |  | > 400 | Yes | Yes |
| **60** |  |  |  | No* |  | > 400 | Yes |  |

For some analogs, we demonstrated here and in previous studies ^22,35^ that virus inhibition did not correlate with inhibition of M2^S31N^ proton currents, even when accounting for slow binding. For instance, compound **38** did not inhibit currents even after 2400 seconds of testing by EP. While it remains possible that a subset of *Amt*(**1**) analogs may possess a limited ability to inhibit M2^S31N^ proton currents, the collective data argue that their primary antiviral targets likely lie elsewhere.

Our studies also indicate that some *Amt*(**1**) analogs do not act by modulating endosomal pH, unlike lysosomotropic agents. Lysosomotropic agents like chlorpromazine, ammonium chloride, chloroquine, and bafilomycin A1 inhibited BPV replication at low or sub-micromolar concentrations, whereas 20 µM of compounds **27** and **38** did not. Notably, this lack of inhibition occurred despite the presence of hydrophobic adducts ^38^ in these analogs, which would be expected to accumulate in intracellular vesicles through membrane permeation by the electroneutral form and increase intravesicular pH.^30^ For example, we recently observed that some IAVs like A/H1N1/PR/8/1934 and A/H1N1/California/07/2009 are inhibited by *Amt*(**1**) analogs bearing different scaffold structures that subtly increase endosomal pH due to their basic character.^19,37^ This increased endosomal pH, in turn, affects HA-mediated fusion, which requires low pH.^,19,39^ While our findings do not rule out this mechanism for all *Amt*(**1**) analogs assessed here, they further support the existence of additional antiviral mechanisms of action. In virus passaging experiments, we observed that M2^WT^ viruses readily developed resistance against *Amt*(**1**) analogs by acquiring V27A or A30T mutations in M2. However, using the A/H1N1/California/07/2009 with M2^S31N^, we showed that resistance to compound **38** occurred between passages 6 and 12. Similarly, the combination of **26**, **27**, and **60** remained effective through 6 passages. Notably, no additional mutations in M2 were detected in these *Amt*(**1**)-resistant viruses, further supporting the presence of other antiviral targets. However, comprehensive virus-wide genome sequencing from additional cultures is needed to identify potential resistance mutations such as those that may arise in HA.^19^ The delayed emergence of resistance in virus with M2^S31N^, compared to the rapid resistance seen in the M2^WT^ virus, suggests that IAV is less capable of overcoming targeting of these alternative antiviral mechanisms. Interestingly, viruses which generated resistance to the combination of compounds **26**, **27**, and **60** retained sensitivity to compound **38**, further supporting that analogs are likely to act on multiple additional antiviral targets. Consequently, the development of *Amt*(**1**)-derived monotherapies or combination therapies optimized against several viral targets, including those identified here (i.e., virus entry and M2-M1 association), reversion of M2^S31N^ to M2^WT^, and potential increases in endosomal pH, may provide a particularly high genetic barrier to future IAV drug resistance.

Notably, different *Amt*(**1**) analogs exhibited a spectrum of overlapping activities. Some were more effective at blocking viral entry, while others excelled at disrupting M2-M1 interactions (**Table 5**). These differences align with our observations that IAV resistant to compounds **26**, **27**, and **60** retain sensitivity to compound **38**. We observed that compound **38** could directly inhibit virus entry, with 50 µM reducing virus entry by ∼50% after just 5 minutes of pre-incubation. Furthermore, compound **38**, unlike compounds **26** and **60**, inhibited H1N1 pseudovirus entry with an EC_50_ of 60 µM. Our previous studies of compound **38** suggest that HA can be a target in some M2^S31N^ viruses such as A/H1N1/California/07/2009.^19^ This may be driven by amino acid variants in HA that affect its binding to cell receptors and/or the efficiency of HA conformational changes at low pH ^19^. It was also reported that A/H1N1/PR/8/1934 is particularly sensitive to increases in endosomal pH caused by the intrinsic basic properties (*i.e.*, alkaline pH-inducing) of *Amt*(**1**) analogs bearing lipophilic scaffold structures.^37^ Conversely, compounds **26** and **60** and others are more effective at disrupting M2-M1 colocalization compared to compound **38**. This is significant as impairment of M1 recruitment to the PM affects viral particle production.^40,41^ However, we have not yet explored whether these *Amt*(**1**) analogs can inhibit other aspects of M1 activity such as during viral entry when M1 dissociates from ribonucleoproteins, allowing them to enter the nucleus. Thus, intraviral buffering that inhibits M1-RNP disruption could be an additional mechanism of action worth investigating.

The structural changes in these *Amt*(**1**)-analogs, though often subtle, resulted in different activities in blocking virus entry and disrupting M2-M1 colocalization. Most *Amt*(**1**) analogs that perturb M2-M1 colocalization have lipophilic adducts attached at the 1- or 2-adamantyl position.

These include 2-(1-adamantyl)piperidine (**26**), 1-adamantyl-1-cyclopentylamine (**27**), 1- adamantyl-1-cyclohexylamine (**28**), 2-n-butyl-2-adamantylamine (**39**) and the spiranic pyrrolidine (**49**) and spirnanic piperidine (**60**). However, while the heterocyclic or carbocyclic amines **26-28, 49, 60,** and **39,** inhibit M2-M1 colocalization, 2-n-propyl-2-adamantylamine (**38**), which also bears an alkyl adduct like **39**, did not. Interestingly, **26-28,** and **39** have higher lipophilicity compared to compound **38** (**Table S2**).

Further support for the relevance of these M2-independent antiviral targets in future therapeutic development comes from a preliminary toxicity study in mice, where compounds **38** and **49** were well tolerated at 30 mg/kg. Although these compounds were more toxic than *Amt*(**1**), which was tolerated at 100 mg/kg, these results support further evaluation of *Amt*(**1**) analogs for absorption, distribution, metabolism, excretion and pharmacokinetics (ADME-PK) as well as *in vivo* efficacy studies.

## 5. Conclusion

In summary, we present a reference study for a series of *Amt*(**1**) variants against various IAV strains and their antiviral mechanisms. We show that some *Amt*(**1**) analogs exhibit low micromolar activities against three M2^S31N^-containing viruses. Specifically, we observed that the 2-propyl-2-adamantyl analog **38** inhibits virus cellular entry, while others like the 1- adamantyl substituted piperidine analog **49** and the 2-adamantyl substituted spiranic pyrrolidine analog **26** disrupt M2-M1 colocalization required for intracellular viral assembly and budding. These patterns of antiviral activities, combined with *in vivo* tolerance of selected analogs, highlight the potential of developing optimized *Amt*(**1**) analogs or combination therapies with higher barriers to drug resistance.

## 6. Abbreviations

ADME-PK: absorption, distribution, metabolism, excretion and pharmacokinetics
Amt: amantadine
EC_50_: half-maximal effective concentration
EP: electrophysiology
HA: hemagglutinin
IAV: influenza A virus
M1: viral matrix protein 1
M2: viral matrix protein 2
PC: positive control
PM: plasma membrane
Rmt: rimantadine

## 7. Acknowledgements

We thank Chiesi Hellas for supporting this research (SARG No 10354), and Professor Nikolas Kolokouris (NKUA) for providing samples of compounds **46** and **47**. The authors thank Dr. Donald Smee for providing Influenza A/H1N1/California/04/2009.

## 8. Author contributions

Conceptualization: A.G., A.K.; Formal analysis: F.B.J., D.C.K.; Funding acquisition: S.C., D.F., I.T.; Investigation: I.V.A., F.B.J., D.C.K., C.M., L.N., A.P., M.R., F.X.S., C.T., I.T., R.Z.; Methodology: S.C., D.F., F.B.J., A.K., D.C.K., L.N., F.X.S.; Supervision: S.C., D.F., A.K., L.N., F.X.S., I.T.; Writing – original draft: A.K., I.T.; Writing – review and editing: S.C., A.K., L.N., A.P., I.T.

## 11. Supporting Information

Detailed Materials and Methods

Table S1, S2

Figure S1-S3

## References

(1) Javanian, M.; Barary, M.; Ghebrehewet, S.; Koppolu, V.; Vasigala, V.; Ebrahimpour, S. A Brief Review of Influenza Virus Infection. J Med Virol 2021, 93 (8), 4638–4646. 10.1002/jmv.26990.

(2) Jones, J. C.; Yen, H.-L.; Adams, P.; Armstrong, K.; Govorkova, E. A. Influenza Antivirals and Their Role in Pandemic Preparedness. Antiviral Res 2023, 210, 105499. 10.1016/j.antiviral.2022.105499.

(3) Jalily, P. H.; Duncan, M. C.; Fedida, D.; Wang, J.; Tietjen, I. Put a Cork in It: Plugging the M2 Viral Ion Channel to Sink Influenza. Antiviral Res 2020, 178, 104780. 10.1016/j.antiviral.2020.104780.

(4) Rossman, J. S.; Jing, X.; Leser, G. P.; Balannik, V.; Pinto, L. H.; Lamb, R. A. Influenza Virus M2 Ion Channel Protein Is Necessary for Filamentous Virion Formation. J Virol 2010, 84 (10), 5078–5088. 10.1128/JVI.00119-10.

(5) Rossman, J. S.; Jing, X.; Leser, G. P.; Lamb, R. A. Influenza Virus M2 Protein Mediates ESCRT-Independent Membrane Scission. Cell 2010, 142 (6), 902–913. 10.1016/j.cell.2010.08.029.

(6) Chen, B. J.; Lamb, R. A. Mechanisms for Enveloped Virus Budding: Can Some Viruses Do without an ESCRT? Virology 2008, 372 (2), 221–232. 10.1016/j.virol.2007.11.008.

(7) Rossman, J. S.; Lamb, R. A. Influenza Virus Assembly and Budding. Virology 2011, 411 (2), 229–236. 10.1016/j.virol.2010.12.003.

(8) Cady, S. D.; Schmidt-Rohr, K.; Wang, J.; Soto, C. S.; DeGrado, W. F.; Hong, M. Structure of the Amantadine Binding Site of Influenza M2 Proton Channels in Lipid Bilayers. Nature 2010, 463 (7281), 689–692. 10.1038/nature08722.

(9) Wright, A. K.; Batsomboon, P.; Dai, J.; Hung, I.; Zhou, H. X.; Dudley, G. B.; Cross, T. A. Differential Binding of Rimantadine Enantiomers to Influenza A M2 Proton Channel. J Am Chem Soc 2016, 138 (5), 1506–1509. 10.1021/jacs.5b13129.

(10) Ma, C.; Polishchuk, A. L.; Ohigashi, Y.; Stouffer, A. L.; Schon, A.; Magavern, E.; Jing, X.; Lear, J. D.; Freire, E.; Lamb, R. A.; DeGrado, W. F.; Pinto, L. H. Identification of the Functional Core of the Influenza A Virus A/M2 Proton-Selective Ion Channel. Proc Natl Acad Sci U S A 2009, 106 (30), 12283–12288. 0905726106 [pii]\r10.1073/pnas.0905726106 [doi].

(11) Thomaston, J. L.; Polizzi, N. F.; Konstantinidi, A.; Wang, J.; Kolocouris, A.; Degrado, W. F. Inhibitors of the M2 Proton Channel Engage and Disrupt Transmembrane Networks of Hydrogen-Bonded Waters. J Am Chem Soc 2018, 140 (45), 15219–15226. 10.1021/jacs.8b06741.

(12) Thomaston, J. L.; Konstantinidi, A.; Liu, L.; Lambrinidis, G.; Tan, J.; Caffrey, M.; Wang, J.; Degrado, W. F.; Kolocouris, A. X-Ray Crystal Structures of the Influenza M2 Proton Channel Drug-Resistant V27A Mutant Bound to a Spiro-Adamantyl Amine Inhibitor Reveal the Mechanism of Adamantane Resistance. Biochemistry 2020, 59 (4), 627–634. 10.1021/acs.biochem.9b00971.

(13) Andreas, L. B.; Barnes, A. B.; Corzilius, B.; Chou, J. J.; Miller, E. A.; Caporini, M.; Rosay, M.; Griffin, R. G. Dynamic Nuclear Polarization Study of Inhibitor Binding to the M2 18–60 Proton Transporter from Influenza A. Biochemistry 2013, 52 (16), 2774–2782. 10.1021/bi400150x.

(14) Dong, G.; Peng, C.; Luo, J.; Wang, C.; Han, L.; Wu, B.; Ji, G.; He, H. Adamantane- Resistant Influenza a Viruses in the World (1902-2013): Frequency and Distribution of M2 Gene Mutations. PLoS One 2015, 10 (3), 1–20. 10.1371/journal.pone.0119115.

(15) Aldrich, P. E.; Hermann, E. C.; Meier, W. E.; Paulshock, M.; Prichard, W. W.; Snyder, J. A.; Watts, J. C. Antiviral Agents. 2. Structure-Activity Relationships of Compounds Related to 1-Adamantanamine. J Med Chem 1971, 14 (6), 535–543. 10.1021/jm00288a019.

(16) Kolocouris, N.; Foscolos, G. B.; Kolocouris, A.; Marakos, P.; Pouli, N.; Fytas, G.; Ikeda, S.; De Clercq, E. Synthesis and Antiviral Activity Evaluation of Some Aminoadamantane Derivatives. J Med Chem 1994, 37 (18), 2896–2902. 10.1021/jm00044a010.

(17) Stamatiou, G.; Foscolos, G. B.; Fytas, G.; Kolocouris, A.; Kolocouris, N.; Pannecouque, C.; Witvrouw, M.; Padalko, E.; Neyts, J.; Clercq, E. D. Heterocyclic Rimantadine Analogues with Antiviral Activity. Bioorg Med Chem 2003, 11 (24). 10.1016/j.bmc.2003.09.024.

(18) Zoidis, G.; Kolocouris, N.; Foscolos, G. B.; Kolocouris, A.; Fytas, G.; Karayannis, P.; Padalko, E.; Neyts, J.; De Clercq, E. Are the 2-Isomers of the Drug Rimantadine Active Anti-Influenza A Agents? Antivir Chem Chemother 2003, 14 (3).

(19) Kolocouris, A.; Tzitzoglaki, C.; Johnson, F. B.; Zell, R.; Wright, A. K.; Cross, T. A.; Tietjen, I.; Fedida, D.; Busath, D. D. Aminoadamantanes with Persistent in Vitro Efficacy against H1N1 (2009) Influenza A. J Med Chem 2014, 57 (11), 4629–4639. 10.1021/jm500598u.

(20) Balannik, V.; Carnevale, V.; Fiorin, G.; Levine, B. G.; Lamb, R. A.; Klein, M. L.; DeGrado, W. F.; Pinto, L. H. Functional Studies and Modeling of Pore-Lining Residue Mutants of the Influenza A Virus M2 Ion Channel. Biochemistry 2010, 49 (4), 696–708. 10.1021/bi901799k.

(21) Wang, J.; Ma, C.; Balannik, V.; Pinto, L. H.; Lamb, R. A.; Degrado, W. F. Exploring the Requirements for the Hydrophobic Scaffold and Polar Amine in Inhibitors of M2 from Influenza A Virus. ACS Med Chem Lett 2011, 2 (4), 307–312. 10.1021/ml100297w.

(22) Drakopoulos, A.; Tzitzoglaki, C.; McGuire, K.; Hoffmann, A.; Ma, C.; Freudenberger, K.; Konstantinidi, A.; Kolocouris, D.; Hutterer, J.; Gauglitz, G.; Wang, J.; Schmidtke, M.; Busath, D. D.; Kolocouris, A.; Kolokouris, D.; Ma, C.; Freudenberger, K.; Hutterer, J.; Gauglitz, G.; Wang, J.; Schmidtke, M.; Busath, D. D.; Kolocouris, A. Unraveling the Binding, Proton Blockage, and Inhibition of Influenza M2 WT and S31N by Rimantadine Variants. ACS Med Chem Lett 2018, 9 (3), 198–203. 10.1021/acsmedchemlett.7b00458.

(23) Stampolaki, Μ.; Hoffmann, A.; Tekwani, K.; Georgiou, K.; Tzitzoglaki, C.; Ma, C.; Becker, S.; Schmerer, P.; Döring, K.; Stylianakis, I.; Turcu, A. L.; Wang, J.; Vázquez, S.; Andreas, L. B.; Schmidtke, M.; Kolocouris, A. A Study of the Activity of Adamantyl Amines against Mutant Influenza A M2 Channels Identified a Polycyclic Cage Amine Triple Blocker, Explored by Molecular Dynamics Simulations and Solid-State NMR**. ChemMedChem 2023. 10.1002/cmdc.202300182.

(24) Chen, H.-W.; Cheng, J. X.; Liu, M.-T.; King, K.; Peng, J.-Y.; Zhang, X.-Q.; Wang, C.-H.; Shresta, S.; Schooley, R. T.; Liu, Y.-T. Inhibitory and Combinatorial Effect of Diphyllin, a v-ATPase Blocker, on Influenza Viruses. Antiviral Res 2013, 99 (3), 371–382. 10.1016/j.antiviral.2013.06.014.

(25) Jang, Y.; Shin, J. S.; Yoon, Y.-S.; Go, Y. Y.; Lee, H. W.; Kwon, O. S.; Park, S.; Park, M.-S.; Kim, M. Salinomycin Inhibits Influenza Virus Infection by Disrupting Endosomal Acidification and Viral Matrix Protein 2 Function. J Virol 2018, 92 (24). 10.1128/JVI.01441-18.

(26) Laporte, M.; Stevaert, A.; Raeymaekers, V.; Boogaerts, T.; Nehlmeier, I.; Chiu, W.; Benkheil, M.; Vanaudenaerde, B.; Pöhlmann, S.; Naesens, L. Hemagglutinin Cleavability, Acid Stability, and Temperature Dependence Optimize Influenza B Virus for Replication in Human Airways. J Virol 2019, 94 (1). 10.1128/JVI.01430-19.

(27) Jalily, P. H.; Eldstrom, J.; Miller, S. C.; Kwan, D. C.; Tai, S. S.-H.; Chou, D.; Niikura, M.; Tietjen, I.; Fedida, D. Mechanisms of Action of Novel Influenza A/M2 Viroporin Inhibitors Derived from Hexamethylene Amiloride. Mol Pharmacol 2016, 90 (2), 80–95. 10.1124/mol.115.102731.

(28) Hintermann, B.; Knupp, M.; Barg, A. Supramalleolar Osteotomies for the Treatment of Ankle Arthritis. J Am Acad Orthop Surg 2016, 24 (7), 424–432. 10.5435/JAAOS-D-12-00124.

(29) Appleyard, G. Amantadine Resistance as a Genetic Marker for Influenza Viruses. Journal of General Virology 1977, 36 (2), 249–255. 10.1099/0022-1317-36-2-249.

(30) Scholtissek, C.; Quack, G.; Klenk, H. D.; Webster, R. G. How to Overcome Resistance of Influenza A Viruses against Adamantane Derivatives. Antiviral Res 1998, 37, 83–95. 10.1016/S0166-3542(97)00061-2.

(31) Petrich, A.; Dunsing, V.; Bobone, S.; Chiantia, S. Influenza A M2 Recruits M1 to the Plasma Membrane: A Fluorescence Fluctuation Microscopy Study. Biophys J 2021, 120 (24), 5478–5490. 10.1016/j.bpj.2021.11.023.

(32) Petrich, A.; Dunsing, V.; Bobone, S.; Chiantia, S. Influenza A M2 Recruits M1 to the Plasma Membrane: A Fluorescence Fluctuation Microscopy Study. Biophys J 2021, 120 (24), 5478–5490. 10.1016/j.bpj.2021.11.023.

(33) Mathiasen, J. R.; Moser, V. C. The Irwin Test and Functional Observational Battery (FOB) for Assessing the Effects of Compounds on Behavior, Physiology, and Safety Pharmacology in Rodents. Curr Protoc Pharmacol 2018, 83 (1), e43. 10.1002/cpph.43.

(34) Torres, E.; Fernández, R.; Miquet, S.; Font-Bardia, M.; Vanderlinden, E.; Naesens, L.; Vázquez, S. Synthesis and Anti-Influenza a Virus Activity of 2,2-Dialkylamantadines and Related Compounds. ACS Med Chem Lett 2012, 3 (12), 1065–1069. 10.1021/ml300279b.

(35) Tzitzoglaki, C.; Wright, A.; Freudenberger, K.; Hoffmann, A.; Tietjen, I.; Stylianakis, I.; Kolarov, F.; Fedida, D.; Schmidtke, M.; Gauglitz, G.; Cross, T. A.; Kolocouris, A. Binding and Proton Blockage by Amantadine Variants of the Influenza M2WT and M2S31N Explained. J Med Chem 2017, 60 (5), 1716–1733. 10.1021/acs.jmedchem.6b01115.

(36) Day, P. M.; Lowy, D. R.; Schiller, J. T. Papillomaviruses Infect Cells via a Clathrin- Dependent Pathway. Virology 2003, 307 (1), 1–11. 10.1016/s0042-6822(02)00143-5.

(37) Torres, E.; Duque, M. D.; Vanderlinden, E.; Ma, C.; Pinto, L. H.; Camps, P.; Froeyen, M.; Vázquez, S.; Naesens, L. Role of the Viral Hemagglutinin in the Anti-Influenza Virus Activity of Newly Synthesized Polycyclic Amine Compounds. Antiviral Res 2013, 99 (3), 281–291. 10.1016/j.antiviral.2013.06.006.

(38) Pisonero-Vaquero, S.; Medina, D. L. Lysosomotropic Drugs: Pharmacological Tools to Study Lysosomal Function. Curr Drug Metab 2017, 18 (12), 1147–1158. 10.2174/1389200218666170925125940.

(39) Daniels, R. Fusion Mutants of the Influenza Virus Hemagglutinin Glycoprotein. Cell 1985, 40 (2), 431–439. 10.1016/0092-8674(85)90157-6.

(40) Wang, D.; Harmon, A.; Jin, J.; Francis, D. H.; Christopher-Hennings, J.; Nelson, E.; Montelaro, R. C.; Li, F. The Lack of an Inherent Membrane Targeting Signal Is Responsible for the Failure of the Matrix (M1) Protein of Influenza A Virus To Bud into Virus-Like Particles. J Virol 2010, 84 (9), 4673–4681. 10.1128/JVI.02306-09.

(41) Liu, H.; Grantham, M. L.; Pekosz, A. Mutations in the Influenza A Virus M1 Protein Enhance Virus Budding To Complement Lethal Mutations in the M2 Cytoplasmic Tail. J Virol 2018, 92 (1). 10.1128/JVI.00858-17.

(42) Chen, B. J.; Leser, G. P.; Jackson, D.; Lamb, R. A. The Influenza Virus M2 Protein Cytoplasmic Tail Interacts with the M1 Protein and Influences Virus Assembly at the Site of Virus Budding. J Virol 2008, 82 (20), 10059–10070. 10.1128/jvi.01184-08.

