## Supplemental Info for "Antiviral Mechanisms and Preclinical Evaluation of Amantadine Analogs that Continue to Inhibit Influenza A Viruses with M2^S31N^-Based Drug Resistance"

**Supporting Information**

Detailed Materials and Methods

Table S1, S2

Figure S1-S3

**Supporting material**

### 2. Materials and Methods

*2.1. Cells, viruses, compounds, and reagents*

Madin-Darby Canine Kidney (MDCK) cells [American Type Culture Collection (ATCC), Kielpin Lomianki, Poland, #CCL-34, NBL-2) were cultured in Dulbecco's Modified Eagle's Medium (DMEM, Sigma-Aldrich, Munich, Germany) supplemented with 0.11% sodium bicarbonate, 5% Cosmic calf serum (Hyclone, Logan, UT, USA), 10 mM (4-(2-hydroxyethyl)-1-piperazineethanesulfonic acid) (HEPES) buffer (Sigma-Aldrich, Munich, Germany), and 50 µg/ml of gentamicin (Sigma-Aldrich, Munich, Germany). For culture of virus stocks and virus infectivity assays, 0.125% bovine serum albumin (BSA, Sigma-Aldrich, Munich, Germany) replaced the Cosmic calf serum. HEK-293T cells (CRL-3216TM, purchased from ATCC, Kielpin Lomianki, Poland) were maintained in phenol red-free, high glucose DMEM plus 2 mM L-glutamine, 100 U/mL penicillin, 100 µg/mL streptomycin and 10% fetal calf serum (FCS) All HEK-293T cell culture products were purchased from PAN-Biotech (Aidenbach, Germany). tsA-201 cells, a derivative of HEK-293T cells (MilliporeSigma Canada, Oakville, ON, Canada), were cultured in modified Eagle's medium + 10% FCS, 100 U/mL penicillin, and 100 µg/mL streptomycin (E10+ medium). All tsA-201 cell culture products were purchased from MilliporeSigma Canada (Oakville, ON, Canada).

The following IAV strains were obtained from ATCC: A/H3N2/Victoria/3/1975 (ATCC VR-822); A/H1N1/PR/8/1934 (ATCC VR-95); A/H1N1/WS/1933 (ATCC VR-1520); and A2/H2N2/Taiwan/1/1964 (ATCC VR-480). The 2009 pandemic strain A/H1N1/California/07/2009 was provided by Dr. Don Smee (Utah State University, Logan, UT, USA).

Murine leukemia virus (MLV)-based pseudovirus encoding firefly luciferase (fLuc) and bearing the HA and neuraminidase (NA) proteins of A/H1N1/Virginia/ATCC3/2009 was produced as we previously described.[26] Briefly, HEK-293T cells were seeded in 6-well plates and transfected with four plasmids encoding MLV gag-pol, the fLuc reporter (kind gift from Stefan Pöhlmann, Göttingen, Germany) and the HA and NA proteins[2626] After 4 hours, the cells were treated with medium containing 2% FCS and incubated for 72 hours at 37 °C. HA was activated by adding 80 µg/mL TPCK-treated trypsin (Sigma-Aldrich, Munich, Germany) and incubating for 15 minutes, followed by 80 µg/mL soybean trypsin inhibitor (Sigma-Aldrich, Munich, Germany). The pseudovirus-containing supernatant was harvested, clarified by centrifugation, and stored at -80˚C.

*2.2. Compounds*

We resynthesized and fully characterized the tested adamantyl amines and analogs as previously reported according to published procedures; **11-19**, [1] **21-23**, [2] **25**, [3,4] **26**, [5] **27**, [6]**28**, [5] **35-43**, [7] **46,** [4] **47,** [4] **49,** [4] **50**, [4] **51**, [8]**52-57**, [9,8] **59**, [10] **60** [5]. To ensure comparability with our previous study, [23] where a larger set of 57 compounds was tested against the A/H1N1/WSN/1933 viruses, the same compound numbering is maintained here.

*2.3. Miniplaque assays*

Assays with MDCK cells were performed as described previously [19]. Briefly, cells were seeded on 12-mm glass cover slips in shell vials and grown until 80-99% confluency. Cultures were washed with serum-free DMEM followed by the addition of 1 mL per vial of media containing 0.125% BSA and test agents at final concentrations of 50, 20, 10, 5, and, if necessary, 2 µM. Stock virus in media with 0.125% BSA was treated with 1.0 µg/mL trypsin for

30 minutes and added to cells, which were then incubated at 33 °C overnight. Following incubation, cultures were washed with phosphate buffered saline (PBS), fixed in -80 °C acetone, and stained with anti-influenza A, FITC-labeled monoclonal antibody (ThermoFisher Scientific, Waltham, MA, USA). $EC_{50}$ determinations were carried out with a fluorescence microscope by counting miniplaques (clusters of infected cells, typically 1-100 per cover slip) in confluent MDCK monolayers. Two to four replicate cultures were included at each drug concentration step. $EC_{50}$s were fitted using the Levenberg-Marquardt algorithm in KaleidaGraph (Synergy Software, Reading, PA, USA).

To assess pre-exposure of virus to drug before cell infection, trypsin-activated virus (A/H1N1/California/07/2009) was generated in MDCK P61 cells and allowed to equilibrate for 30 minutes at room temperature with a 1:8 virus dilution in DMEM containing 0.125% BSA. The mixture was then split into two cultures, 1 mL each: one for control (no drug) and one for drug **38** at 50 μM. 10 μL samples were taken at various times post drug addition and cultured by miniplaque assay in duplicate.

*2.4. M2 electrophysiology*

A pcDNA3 plasmid encoding full-length $M2^{S31N}$ (A/H1N1/California/07/2009) was co-transfected with pcDNA3 plasmid encoding green fluorescent protein (GFP) into tsA-201 cells as described previously.[27] After 24 hours, single GFP-positive cells were perfused continuously at 3-5 mL/minutes with bath solution containing 150 mM N-methyl-D-glucamine (NMDG), 10 mM HEPES, 10 mM D-glucose, 2 mM $CaCl_2$, and 1 mM $MgCl_2$ buffered at pH 7.4. For pH 5.5 solution, HEPES was replaced by 4-morpholineethanesulfonic acid (MES). Patch electrodes were pulled from thin-walled borosilicate glass (World Precision Instruments, Sarasota, FL) and fire polished before filling with standard pipette solution containing 140 mM NMDG, 10 mM egtazic acid (EGTA), 10 mM HEPES, and 1 mM $MgCl_2$ buffered at pH 7.2. Voltage clamp experiments were performed with an Axopatch 200B amplifier (Molecular Devices, Sunnyvale, CA) connected to a Digidata1322A 16-bit digitizer as previously described.[28] All cell culture reagents were purchased from MilliporeSigma Canada (Oakville, ON, Canada).

*2.5. Resistance test plaque sequencing*

MDCK cells were incubated with IAV (at a multiplicity of infection (MOI) of 1) for one hour to allow virus adsorption. The excess virus was then washed off PBS, and the cells were incubated with Eagle's minimal essential medium supplemented with FBS and the test agent for 3-4 days. If no cytopathic effect was observed, 0.5 mL of the supernatant was centrifuged to remove cell detritus and transferred to Petri dishes with confluent MDCK monolayers. If a cytopathic effect was visible, 1 mL of the supernatant was additionally stored at -80 °C. Cells were iteratively incubated for up to 4 days and up to 10 passages. Ten-fold serial dilutions of the viral stocks from passages 1, 4, and 9 were used for plaque assays and viral sequencing as described previously.[29,30]

*2.6. Pseudovirus entry assay*

MDCK cells were seeded in white half-area 96-well plates (Greiner Bio-One, Vilvoorde, Belgium) and incubated for 24 hours. The cells were then preincubated for 20 minutes with serial dilutions of the compounds, followed by exposure to MLV-based pseudovirus and the compounds for 2 hours. After the removal of excess virus and compound, cells were incubated for 72 hours. fLuc activity was measured using the luciferase assay system kit and GloMax Navigator luminometer (Promega, Madison, WI, USA).

*2.7. Confocal microscopy analysis of M1 intracellular localization*

The plasmid encoding mCherry2 linked to a myristoylated and palmitoylated peptide (mp-mCherry2) was previously described.[31] The plasmid encoding M1 from A/H1N1/WSN/1933 with mEGFP C-terminal fusion (WSN-M1-mEGFP) was a kind gift of Andreas Herrmann (Humboldt University Berlin, Berlin, Germany). The plasmid encoding M2 from A/H1N1/WSN/1933 with RFP C-terminal fusion (WSN-M2-RFP) was kindly provided by Michael Veit (Free University, Berlin, Germany). The plasmid encoding M2 from A/H7N1/FPV/Rostock/1934 with HAsp-mCherry2 N-terminal fusion (HAsp-mCherry2-FPV-M2) was previously described.[31] The HAsp-mCherry2-WSN-M2 construct was generated by amplifying the M2 sequence from WSN-M2-RFP and cloning it into HAsp-mCherry2-FPV-M2 via restriction with XbaI and NotI using standard cloning procedures. [31]

For imaging experiments, 35 mm dishes (CellVis, Mountain View, CA, USA) with an optical glass bottom (#1.5 glass, 0.16–0.19 mm) were coated with 0.01% (w/v) poly-L-lysine (molecular weight [MW] 150,000–300,000 Da, Sigma-Aldrich, Munich, Germany) for 4 hours at 37 °C, rinsed three times with Dulbecco's PBS with $Ca^{2+}$ and $Mg^{2+}$ (DPBS+/+; PAN-Biotech, Aidenbach, Germany), and seeded at $6 \times 10^5$ cells per dish. After 24 h, transfection was performed using Turbofect® (Thermo Fisher Scientific, Waltham, MA, USA) using 100 ng pDNA WSN-M1-mEGFP and 200 ng pDNA HAsp-mCherry2-WSN-M2 or mp-mCherry2, and cells were incubated for a further 24 hours in the presence of *Amt*(**1**) analogs (30 μM) or the analog solvent ($H_2O$). Confocal microscopy was performed on a Zeiss LSM780 system (Carl Zeiss, Oberkochen, Germany). Three to four tile scan images were randomly chosen and automatically acquired per sample, spaced in a regular 3 x 3 pattern, with an image size of 637.23 x 637.23 μm. The percentage of cells displaying M1 signal at the plasma membrane (PM) was determined by manually identifying cells with M1 signal at the PM and calculating the ratio of these cells to the total number of cells in each image using Fiji ImageJ software (**Figure S2**).[32] The relative amount of cells showing M1 membrane localization was normalized to the median value of the positive control (co-expression of WSN-M1-mEGFP and HAsp-mCherry2-FPV-M2). Data from three independent experiments were analyzed using GraphPad Prism vs. 9.0.0 (GraphPad Software, LCC, San Diego, CA, USA). Descriptive statistics are summarized in **Table S1**. Statistical significance was tested by using D'Agostino-Pearson normality test followed by one-way ANOVA analysis and Dunnett's multiple comparisons test.

#### *2.8. Toxicity and Neurotoxicity Testing*

Acute toxicity was assessed in CD-1 mice (Charles River Laboratoires, France) at doses of 30, 100 or 300 mg/kg body weight, administered intraperitoneally (*i.p.*). Mice were 6-10 weeks of age with both sexes used. Experiments were conducted following the Federation of European Laboratory Animal Science Association guidelines, according to European directive 2010/63/EU and under supervision of the Ethical Committee of the Rovira i Virgili University (#T9900003). Animals were housed in standard conditions with free access to food and water and an artificial 12 h light/dark cycle. Neurotoxicity was evaluated using a modified Functional Observational Battery (FOB) at 30, 60 and 120 minutes after *i.p.* administration.[33] Mice were monitored for 48 hours to determine lethality. Behavioral patterns such as rearings, climbing, grooming, defecation and urination were also recorded.

**Table S1.** Descriptive statistics of the image analysis regarding M1 localization at the PM. Data correspond to **Figure 8** of the main manuscript. $N_{cells}$: number of analyzed cells, iqr: interquartile range, sd: standard deviation, sem: standard error of the mean, NC: negative control (WSN-M1-mEGFP + mp-mCherry2), PC: positive control (WSN-M1-mEGFP + $HA_{sp}$-mCherry2-WSN-M2).

| **treatment** | **$N_{cells}$** | **median** | **iqr** | **mean** | **sd** | **sem** |
|---|---|---|---|---|---|---|
| **NC** | 805 | 0.0 | 0.0 | 0.0 | 0.0 | 0.0 |
| **PC** | 1546 | 97.8 | 14.9 | 99.4 | 10.9 | 3.4 |
| **38** | 1880 | 89.7 | 37.3 | 82.7 | 22.6 | 6.8 |
| **49** | 1803 | 53.2 | 21.7 | 55.1 | 14.4 | 4.4 |
| **60** | 1900 | 67.6 | 33.7 | 71.2 | 26.8 | 8.1 |
| **26** | 1790 | 61.0 | 20.9 | 58.7 | 12.9 | 3.9 |
| **28** | 1866 | 52.6 | 27.1 | 57.5 | 24.2 | 7.3 |
| **39** | 1791 | 36.7 | 7.9 | 35.9 | 7.2 | 2.2 |

**Table S2.** Calculated lipophilicity for selected compounds.

| Compound | logP |
|---|---|
| **26** | 4.31 ± 0.27 |
| **27** | 4.25 ± 0.26 |
| **28** | 4.82 ± 0.26 |
| **38** | 3.84 ± 0.24 |
| **39** | 4.37 ± 0.24 |
| **49** | 3.28 ± 0.36 |
| **60** | 3.85 ± 0.36 |

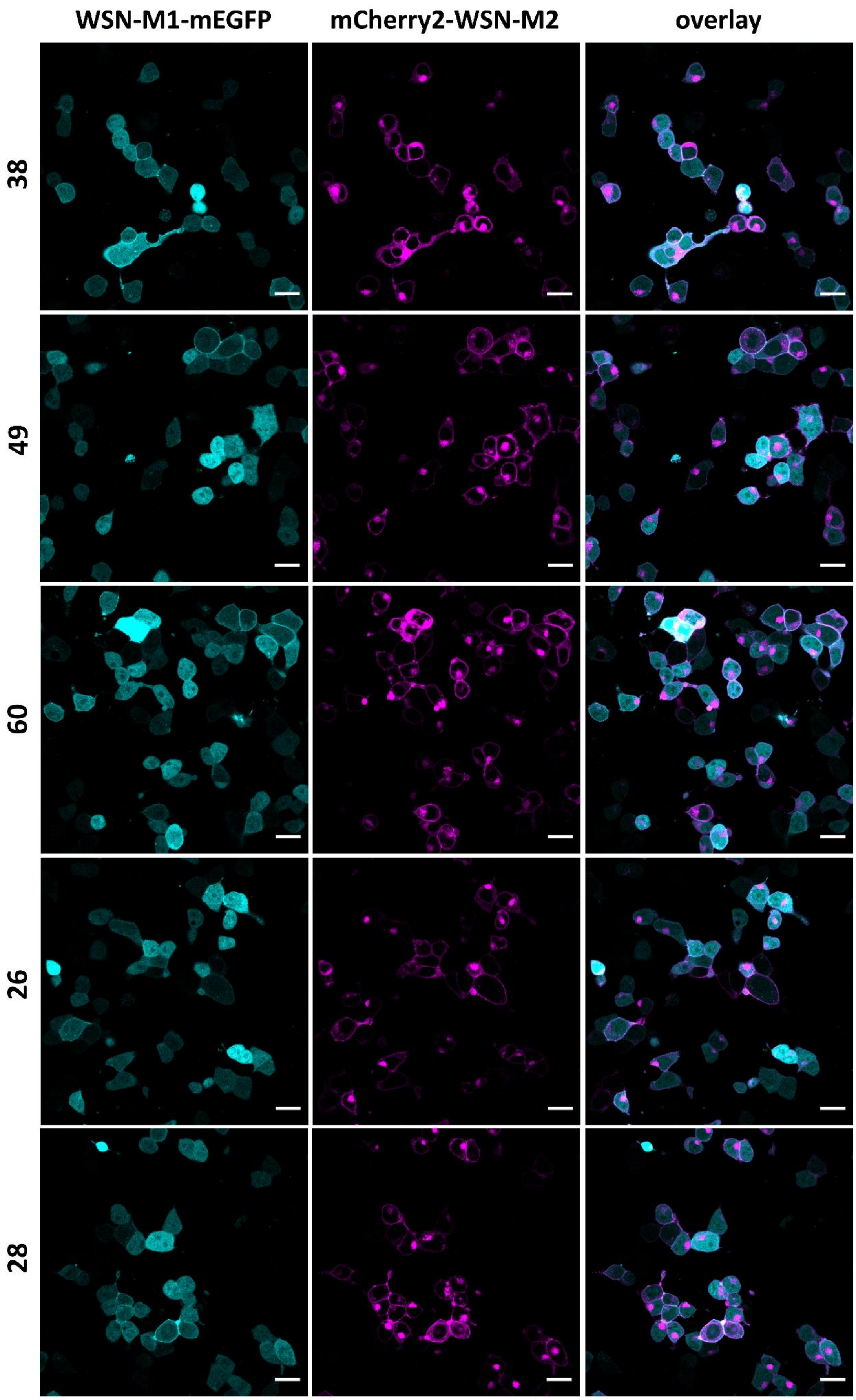
WSN-M1-mEGFP
mCherry2-WSN-M2
overlay
38
49
60
26
28

**Figure S1.** Representative confocal fluorescence images of HEK-293T cells co-expressing WSN-M1-mEGFP (cyan) and $HA_{sp}$-mCherry2-WSN-M2 (magenta), after treatment with different *Amt*(**1**) analogs (30 μM) for 24 hours. The panels on the right show the two channels merged in a single image. Scale bars represent 20 μm.

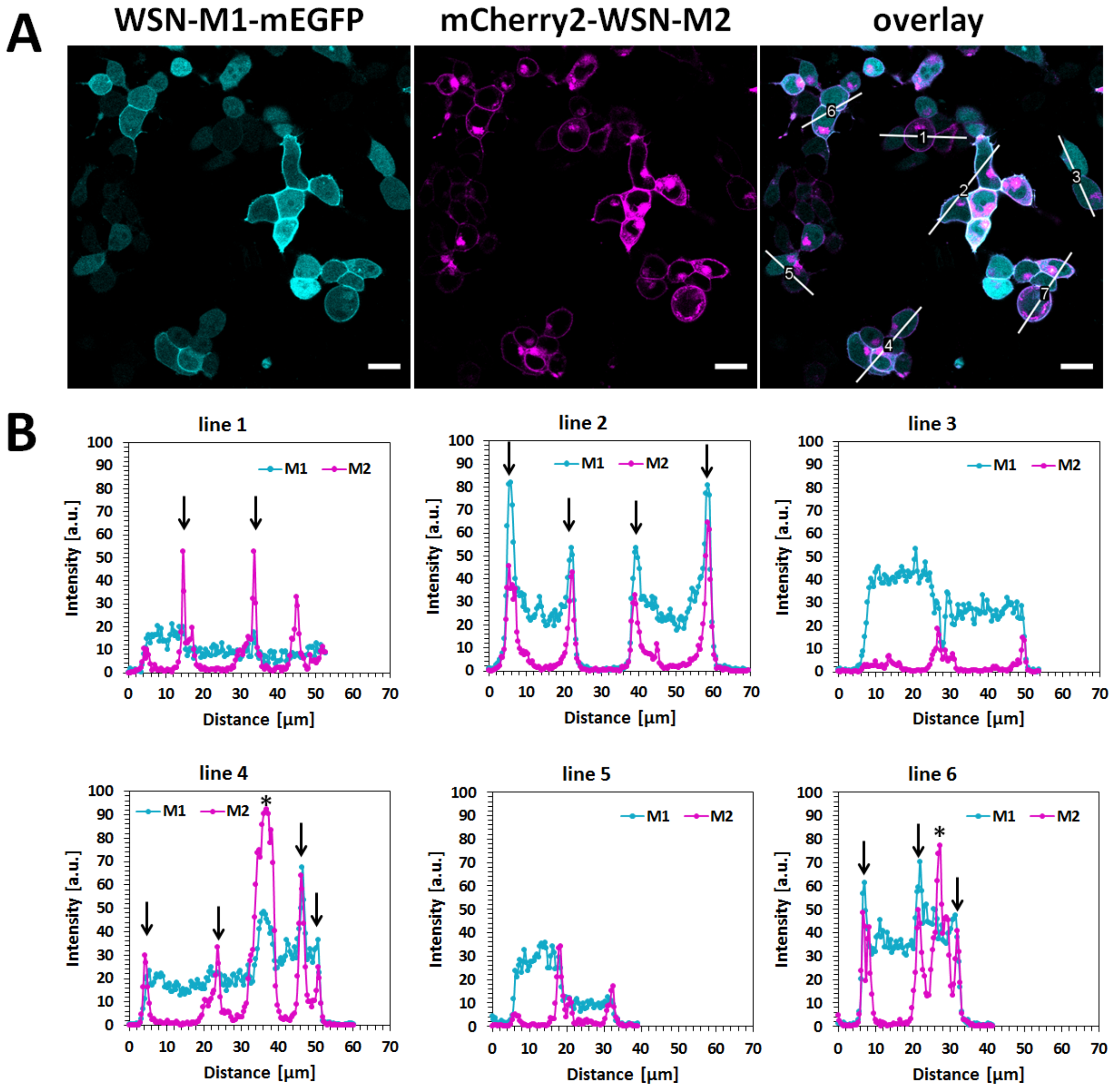


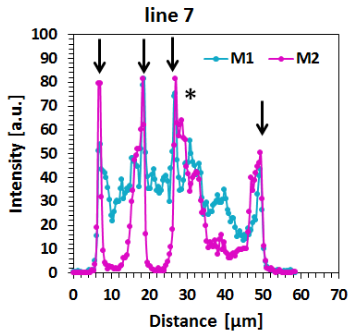


C

| # Line | # M1 PM local. | # total |
|---|---|---|
| 1 | 1 | 3 |
| 2 | 2 | 2 |
| 3 | 0 | 2 |
| 4 | 2 | 4 |
| 5 | 0 | 2 |
| 6 | 2 | 2 |
| 7 | 3 | 3 |

**Figure S2.** Representative analysis for the quantification of M1 plasma membrane localization. **A**, Representative confocal fluorescence images of HEK-293T cells co-expressing WSN-M1-mEGFP (cyan) and $HA_{sp}$-mCherry2-WSN-M2 M2 (magenta). The panel on the right shows the two channels merged in a single image. Plasma membrane localization of M1 was determined by line plot profile analysis of the fluorescence signal of WSN-M1-mEGFP (cyan) and $HA_{sp}$-mCherry2-WSN-M2 M2 (magenta) performed at the indicated solid white lines (indicated here as examples) in the overlay images. The line profiles are shown in (B). Scale bars represent 20 µm. **B**, Exemplar line plot profile analysis of WSN-M1-mEGFP (cyan) and $HA_{sp}$-mCherry2-WSN-M2 M2 (magenta) of the indicated lines in the overlay image shown in (A). Peaks in the magenta lines (i.e., related to M2 distribution along the lines) indicate enrichment at the PM. Arrows illustrate examples of overlap between peaks in cyan and magenta lines. For example, the two arrows indicated for “line 1” refer to the edges of one cell in which M1 partitions at the PM. The four arrows indicated for “line 2” indicate the edges of two cells in which M1 partitions at the PM. For the case of “line 3”, no peak in the cyan line is observed in correspondence with the magenta peak. This indicates lack of visible partition of M1 at the PM of the two cells crossed by “line 3”. The symbol * indicates signal from occasional intracellular accumulation of M2. **C**, Overview of the number of cells with M1 at the plasma membrane (# M1 PM local.) and the total number of cells (# total) per line plot analysis.

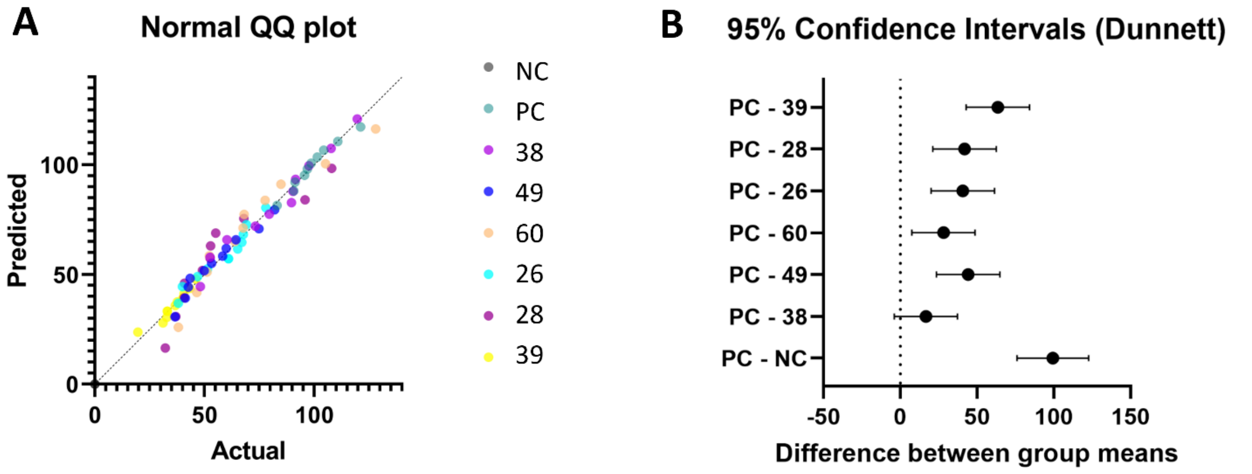


**Figure S3.** Statistical analysis of the PM recruitment of M1 from confocal imaging of three independent experiments. Data corresponds to **Figure 8** of the main manuscript. **A**, Normality test visualized as QQ plot. **B**, Representation of the 95% confidence intervals from the Dunnett's multiple comparison test. NC: negative control (WSN-M1-mEGFP + mp-mCherry2), PC: positive control (WSN-M1-mEGFP + $HA_{sp}$-mCherry2-WSN-M2).

### Supporting information references


(1) Stampolaki, M.; Hoffmann, A.; Tekwani, K.; Georgiou, K.; Tzitzoglaki, C.; Ma, C.; Becker, S.; Schmerer, P.; Döring, K.; Stylianakis, I.; Turcu, A. L.; Wang, J.; Vázquez, S.; Andreas, L. B.; Schmidtke, M.; Kolocouris, A. A Study of the Activity of Adamantyl Amines against Mutant Influenza A M2 Channels Identified a Polycyclic Cage Amine Triple Blocker, Explored by Molecular Dynamics Simulations and Solid-State NMR**. *ChemMedChem* **2023**. https://doi.org/10.1002/cmdc.202300182.

(2) Drakopoulos, A.; Tzitzoglaki, C.; McGuire, K.; Hoffmann, A.; Ma, C.; Freudenberger, K.; Konstantinidi, A.; Kolocouris, D.; Hutterer, J.; Gauglitz, G.; Wang, J.; Schmidtke, M.; Busath, D. D.; Kolocouris, A.; Kolokouris, D.; Ma, C.; Freudenberger, K.; Hutterer, J.; Gauglitz, G.; Wang, J.; Schmidtke, M.; Busath, D. D.; Kolocouris, A. Unraveling the Binding, Proton Blockage, and Inhibition of Influenza M2 WT and S31N by Rimantadine Variants. *ACS Med Chem Lett* **2018**, *9* (3), 198–203. https://doi.org/10.1021/acsmedchemlett.7b00458.

(3) Kolocouris, N.; Kolocouris, A.; Foscolos, G. B.; Fytas, G.; Neyts, J.; Padalko, E.; Balzarini, J.; Snoeck, R.; Andrei, G.; De Clercq, E. Synthesis and Antiviral Activity Evaluation of Some New Aminoadamantane Derivatives. 2. *J Med Chem* **1996**, *39* (17), 3307–3318. https://doi.org/10.1021/jm950891z.

(4) Kolocouris, N.; Foscolos, G. B.; Kolocouris, A.; Marakos, P.; Pouli, N.; Fytas, G.; Ikeda, S.; De Clercq, E. Synthesis and Antiviral Activity Evaluation of Some Aminoadamantane Derivatives. *J Med Chem* **1994**, *37* (18), 2896–2902. https://doi.org/10.1021/jm00044a010.

(5) Stamatiou, G.; Foscolos, G. B.; Fytas, G.; Kolocouris, A.; Kolocouris, N.; Pannecouque, C.; Witvrouw, M.; Padalko, E.; Neyts, J.; Clercq, E. D. Heterocyclic Rimantadine Analogues with Antiviral Activity. *Bioorg Med Chem* **2003**, *11* (24). https://doi.org/10.1016/j.bmc.2003.09.024.

(6) Tzitzoglaki, C.; Drakopoulos, A.; Konstantinidi, A.; Stylianakis, I.; Stampolaki, M.; Kolocouris, A. Approaches to Primary Tert-Alkyl Amines as Building Blocks. *Tetrahedron* **2019**, *75* (34), 130408. https://doi.org/10.1016/j.tet.2019.06.016.

(7) Kolocouris, A.; Tzitzoglaki, C.; Johnson, F. B.; Zell, R.; Wright, A. K.; Cross, T. A.; Tietjen, I.; Fedida, D.; Busath, D. D. Aminoadamantanes with Persistent in Vitro Efficacy against H1N1 (2009) Influenza A. *J Med Chem* **2014**, *57* (11), 4629–4639. https://doi.org/10.1021/jm500598u.

(8) Georgiou, K.; Konstantinidi, A.; Hutterer, J.; Freudenberger, K.; Kolarov, F.; Lambrinidis, G.; Stylianakis, I.; Stampelou, M.; Gauglitz, G.; Kolocouris, A. Accurate Calculation of Affinity Changes to the Close State of Influenza A M2 Transmembrane Domain in Response to Subtle Structural Changes of Adamantyl Amines Using Free Energy Perturbation Methods in Different Lipid Bilayers. *Biochimica et Biophysica Acta (BBA) - Biomembranes* **2024**, *1866* (2), 184258. https://doi.org/10.1016/j.bbamem.2023.184258.

(9) Stylianakis, I.; Kolocouris, A.; Kolocouris, N.; Fytas, G.; Foscolos, G. B.; Padalko, E.; Neyts, J.; De Clercq, E. Spiro[Pyrrolidine-2,2′-Adamantanes]: Synthesis, Anti-Influenza Virus Activity and Conformational Properties. *Bioorg Med Chem Lett* **2003**, *13* (10), 1699–1703. https://doi.org/10.1016/S0960-894X(03)00231-2.

(10) Tzitzoglaki, C.; Wright, A.; Freudenberger, K.; Hoffmann, A.; Tietjen, I.; Stylianakis, I.; Kolarov, F.; Fedida, D.; Schmidtke, M.; Gauglitz, G.; Cross, T. A.; Kolocouris, A. Binding and Proton Blockage by Amantadine

Variants of the Influenza M2WT and M2S31N Explained. *J Med Chem* **2017**, *60* (5), 1716–1733. https://doi.org/10.1021/ACS.JMEDCHEM.6B01115/SUPPL_FILE/JM6B01115_SI_002.CSV.
